# *In vivo* analysis of the relationship between CP26 and qE-type NPQ via higher-order *Arabidopsis cp26* mutants

**DOI:** 10.1101/2024.07.26.605339

**Authors:** Julia Walter, Dhruv Patel-Tupper, Lam Lam, Alexa Ma, Georgia Taylor, Graham R. Fleming, Krishna K. Niyogi, Johannes Kromdijk

## Abstract

CP26 is a monomeric minor light-harvesting complex of PSII (LHCII) protein that connects major LHCII trimers to the PSII core in photosynthetic thylakoid membranes. Previous studies have proposed that CP26 is not only involved in light harvesting but could also be involved in non-photochemical quenching (NPQ). Here, we analyzed higher-order *Arabidopsis cp26* mutants using biophysical and pharmacological approaches to investigate the nature of NPQ and its relationship to known NPQ regulators (PSII subunit S (PsbS), the xanthophyll-converting enzyme VDE and the pH gradient across the thylakoid membrane). Maximum PSII quantum efficiencies (F_v_/F_m_) and chlorophyll fluorescence lifetimes in the dark were significantly lower in *cp26* mutants, confirming that CP26 deficiency leads to a sustained quenched state even in the absence of light. Destabilized PSII-LHCII supercomplexes as observed with native PAGE analysis are the likely cause for this pre-quenched state, without other antenna proteins being able to rescue this phenotype. Further analyses revealed that *cp26* mutants exhibit modest (single mutant) to highly significant (double mutants) reductions in overall NPQ capacity, which do not directly rely on PsbS and VDE (although the effect is more pronounced when these qE components are altered) but depend on thylakoid lumen acidification and protonation of protein residues. Together, these results show that the NPQ component lacking in *cp26* mutants acts independently of qE and qZ and is induced in a slower phase of NPQ induction that most likely relies on pH-dependent conformational changes.

## Introduction

Under sub-optimal environmental conditions, such as high light and/or cold stress, photoprotective mechanisms protect the photosynthetic electron transport chain from photooxidative damage by preventing the formation of harmful reactive oxygen species. The dissipation of excess light energy as heat is commonly referred to as non-photochemical quenching (NPQ) (for recent reviews see Bassi and Dall’Osto, 2021; Ruban and Wilson, 2021). The fastest NPQ component is termed energy-dependent quenching (qE) and, in plants, relies on the Photosystem II (PSII) subunit S (PsbS) as a lumenal pH sensor (Li *et al*., 2000, 2002b; Nicol and Croce, 2021). Upon light exposure, lumen-exposed glutamate residues in PsbS are protonated (Li *et al*., 2002a, 2004; Liguori *et al*., 2019), inducing conformational changes and monomerization of PsbS dimers (Bergantino *et al*., 2003; Krishnan *et al*., 2017). PsbS monomers have been proposed to then intercalate into the light-harvesting complexes around PSII (LHCII antenna system) and detach the loosely (L) and moderately (M) bound LHCII trimers from the PSII-LHCII supercomplexes, thereby rearranging the major antennae into detached LHCII aggregates (Kiss *et al*., 2008; Betterle *et al*., 2009; Wilk *et al*., 2013; Ware *et al*., 2015; Correa-Galvis *et al*., 2016; Sacharz *et al*., 2017; Daskalakis *et al*., 2019; Pawlak *et al*., 2020; Nicol and Croce, 2021). The main qE quenching site Q1 has been hypothesized to be located in detached LHCII aggregates (Horton *et al*., 1991; Mullineaux *et al*., 1993; Miloslavina *et al*., 2008, 2011; Goral *et al*., 2012; Tian *et al*. 2015; Natali *et al*., 2016; Adams *et al*., 2018; Shukla *et al*., 2020; Tutkus *et al*., 2021; Wilson *et al*., 2022), however, the actual role of PsbS in qE remains to be determined (reviewed in Marulanda Valencia and Pandit, 2024). In response to decreased lumen pH, the reversible xanthophyll cycle is activated through protonation of the enzyme violaxanthin de-epoxidase (VDE), which converts violaxanthin into antheraxanthin and zeaxanthin upon high light exposure, thereby enhancing qE. Aside from the putative quenching site in trimeric LHCII, zeaxanthin and the xanthophyll lutein bound to the minor antennae appear to form transient radical cations, which could potentially be an independent NPQ mechanism (Holt *et al*., 2005; Ahn *et al*., 2008; Avenson *et al*., 2008; Li *et al*., 2009; Pinnola *et al*., 2016; Leuenberger *et al*., 2017).

A second quenching site, Q2, is proposed in the strongly bound S-LHCII trimers attached to PSII and possibly in the minor antennae (CP29, CP26, CP24), which connect the major LHCII trimers to the PSII core. NPQ occurring in the Q2 site could account for the slower phase of zeaxanthin-dependent NPQ induction (qZ, several minutes to tens of minutes timescale), relying solely on the activity of the xanthophyll cycle (Miloslavina *et al*., 2008, Nilkens *et al*., 2010), although this mechanism has been debated (Saccon *et al*., 2020). *In vitro* reconstitution of recombinant LHCII proteins with the xanthophyll pigments revealed that CP26 binds zeaxanthin much more efficiently than any other LHCII protein (Morosinotto *et al*., 2002). Moreover, CP26 is the only minor antenna protein that has been shown to undergo a conformational change upon binding of zeaxanthin - independently from the presence of PsbS - resulting in a shift in isoelectric point (pI) and reduced fluorescence levels (Dall’Osto *et al*., 2005). *In vitro* chlorophyll-binding site analyses also suggested that CP26 could be involved in quenching (Ballottari *et al*., 2009, 2013).

Here, we investigated the contribution of the minor antenna protein CP26 to NPQ and its relationship to different qE components *in vivo* by studying the single knockout mutant *cp26*, as well as double mutants with modified PsbS (*cp26 npq4* and *cp26 L17*) and VDE (*cp26 npq1*) levels. We found that *cp26* mutants had disrupted PSII-LHCII supercomplexes and maintained a quenched state after dark acclimation. Under high light, however, *cp26* mutants displayed lower NPQ levels than their controls, particularly during the later phase of NPQ induction, confirming that CP26 contributes to a slow form of NPQ. Consistent with previous *in vitro* analyses, the effect of CP26 knockout on NPQ persisted in the absence of PsbS and VDE. However, the effect was removed by inhibition of lumen acidification or blocking of protonatable residues (possibly in CP26 itself). Altogether, these results suggest that in addition to its role in energy transfer, CP26 contributes to a minor NPQ component, which is distinct from qE.

## Results

### *Arabidopsis* mutants lacking CP26 have lower PSII efficiencies in the absence of light due to quenching in the dark

To investigate the relationship between CP26 and the qE components PsbS and VDE, the *Arabidopsis cp26* T-DNA insertion line SALK-014869C was crossed with the PsbS knockout line *npq4*, the PsbS overexpression line L17 and the VDE knockout line *npq1* (for molecular confirmation of double mutants see Fig. S1).

**Fig. S1:**
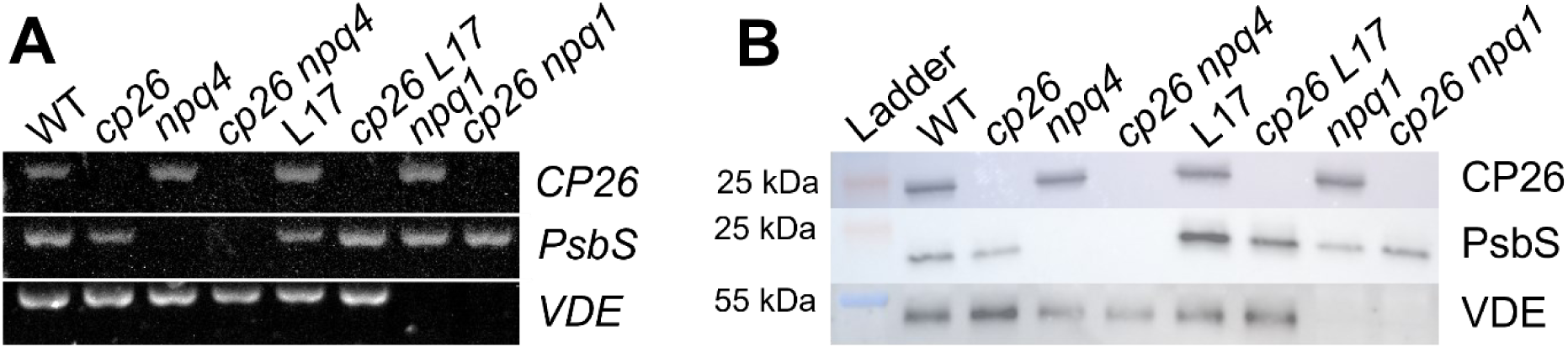
Molecular confirmation of *Arabidopsis* mutants used in this study. **A)** DNA confirmation with PCR of *CP26*, Photosystem II Subunit S (*PsbS*) and violaxanthin de-epoxidase (*VDE*) genes. **B)** Western blot results using CP26, PsbS and VDE antibodies (1 µg chlorophyll of thylakoid membrane protein extracts loaded).

Using biophysical and biochemical assays, the *cp26* mutants were assessed for their photosynthetic and photoprotective capacities in comparison to their respective WT/single mutant controls.

The maximum quantum efficiency of PSII (F_v_/F_m_) was determined after 75 min of dark acclimation by measuring pulse-amplitude-modulated (PAM) chlorophyll fluorescence from intact leaves using a Dual-Klas-NIR/GFS-3000/LI-6800 setup for controlled environmental conditions (temperature, humidity and CO_2_). All *cp26* mutants showed significantly lower F_v_/F_m_ values (∼0.77 ± 0.002-0.005; p<0.0001-0.001) compared to their controls (∼0.80 ± 0.001-0.003; Fig. 1A). A decrease in F_v_/F_m_ either suggests an increase in the minimum fluorescence (F_o_) or a lower dark-acclimated maximum fluorescence (F_m_). While F_o_ levels were slightly but not significantly elevated (apart from *cp26 L17*; p<0.05; Fig. 1B), F_m_ values were significantly lower in all *cp26* single and double mutants compared to their respective controls (Fig. 1C), indicating a quenched state even in the absence of light. Time-correlated single photon counting (TCSPC) measurements on intact leaves also corroborated these data by revealing a decrease in chlorophyll fluorescence lifetime by approximately 0.1 ns in the absence of CP26 across all genotypic pairs (Fig. 1D).

**Fig. 1:**
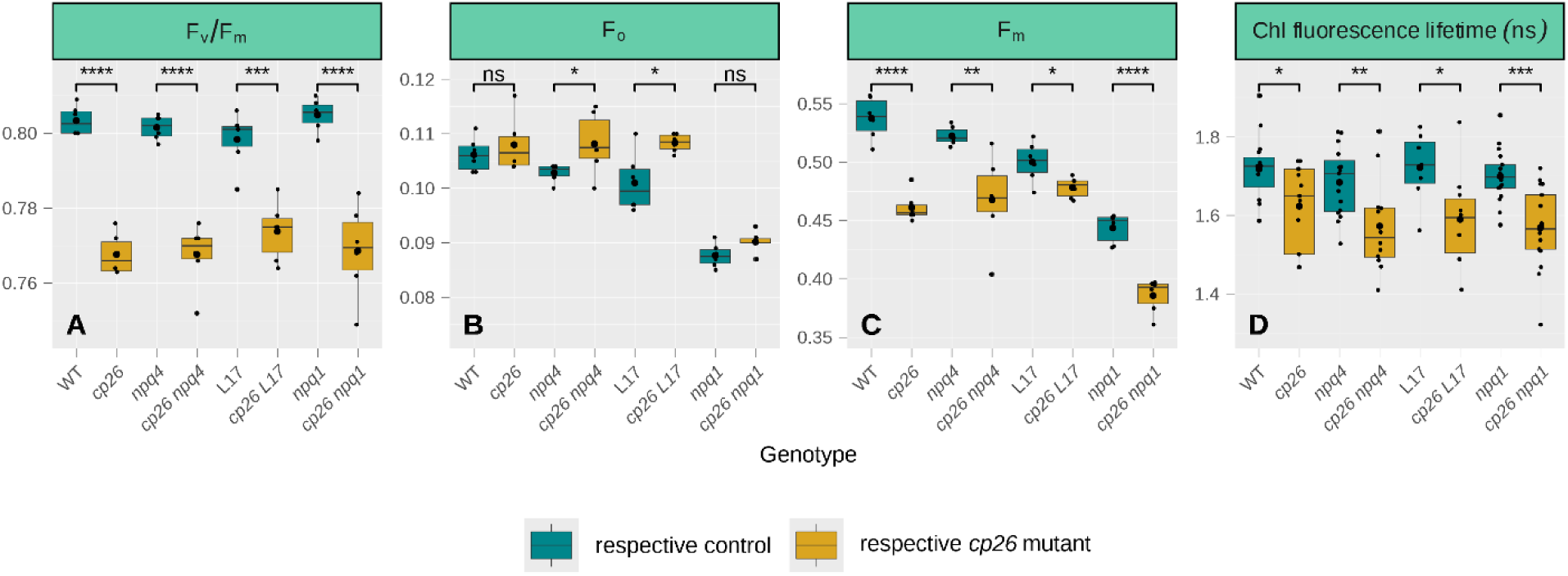
Dark-acclimated chlorophyll fluorescence parameters of *cp26* mutants. **A)** Maximum quantum yield of PSII, F_v_/F_m_, calculated from minimum (**B** F_o_) and maximum (**C** F_m_) chlorophyll fluorescence levels. **D)** Chlorophyll fluorescence lifetime values (in nanoseconds) of intact leaves following dark acclimation. Data were collected from six biological replicates (n=6) for PAM chlorophyll fluorescence measurements and 7-17 biological replicates (n=7-17) for chlorophyll fluorescence lifetime measurements. Boxplots contain individual data points (black dots), the median (black line inside the box) and the mean (larger black dot inside the box). Significant differences between control genotypes (WT, *npq4*, L17 and *npq1*; green boxes) and *cp26* mutants (*cp26*, *cp26 npq4*, *cp26 L17* and *cp26 npq1*; yellow boxes) are indicated by asterisks (Student’s t-test). Significance levels: ns = no significance, * = p<0.05, ** = p<0.01, *** = p<0.001 and **** = p<0.0001.

Following F_v_/F_m_ measurements, chlorophyll fluorescence was recorded during two consecutive cycles of high light (1000 µmol photons m^-2^ s^-1^ for 20 min) and dark (10 min). PSII efficiencies during light phases (F_q_’/F_m_’) were similar between all genotypes, with small differences only observed in the comparison between L17 and *cp26 L17* double mutant (Fig. S2B, bottom panel). In contrast, in the dark phases, most CP26-deficient plants had significantly lower F_v_’/F_m_’ values (PSII efficiencies during dark phases; equivalent to F_q_’/F_m_’ in the light) than their respective controls, except *cp26 npq1* (Fig. S2), and PSII quantum efficiencies were significantly decreased in the second dark phase compared to the first.

**Fig. S2:**
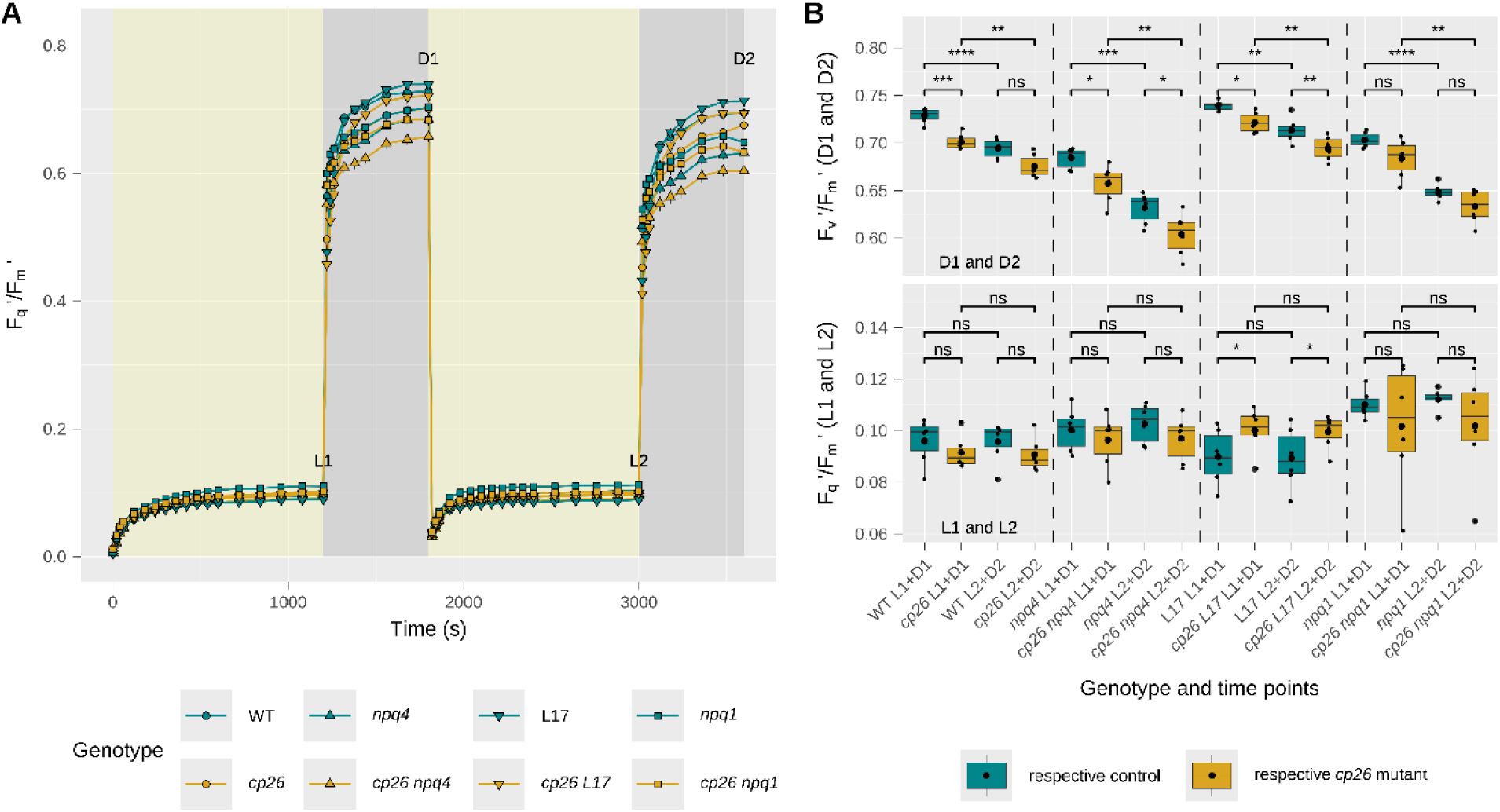
PSII quantum efficiencies in *cp26* and double mutants with contrasting PsbS (knockout = *npq4* and overexpression = L17) and VDE (*npq1*) alleles. **A)** PSII quantum yields, F_q_’/F_m_’, measured upon high light induction (1000 µmol photons m^-2^ s^-1^ for 20 min, yellow background), followed by 10 min of dark relaxation (dark grey background), in two consecutive cycles. **B)** Maximum F_v_’/F_m_’ values (top panel, measured at the end of the two dark phases and annotated as D1 and D2) and maximum F_q_’/F_m_’ values (bottom panel, measured at the end of the two high light phases and annotated as L1 and L2). Line plots show mean values and error bars indicate the standard error of the mean for control genotypes (WT, *npq4*, L17 and *npq1*; green lines and circles, upwards-pointing triangles, downwards-pointing triangles, and squares, respectively) and *cp26* mutants (*cp26*, *cp26 npq4*, *cp26 L17* and *cp26 npq1*; yellow lines and circles, upwards-pointing triangles, downwards-pointing triangles, and squares, respectively). Boxplots contain individual data points (black dots), the median (black line inside the box) and the mean (larger black dot inside the box). Data were collected from six biological replicates (n=6). Significant differences between control genotypes (green boxes) and *cp26* mutants (yellow boxes) are indicated by asterisks (two-way repeated measures ANOVA with following paired t-test as post-hoc test within D1 & D2 and L1 & L2 maximum values). To compare D1 & D2 and L1 & L2 time points within genotypes, a non-paired Student’s t-test was applied. Significance levels: * = p<0.05, ** = p<0.01, *** = p<0.001 and **** = p<0.0001.

### Absence of CP26 affects NPQ and PSII quantum efficiencies during illumination due to impairments in PSII-LHCII supercomplex stability

NPQ was calculated from the same two cycles of 20 min high light – 10 min darkness (Fig. 2A). During high light exposure, NPQ in WT was rapidly induced during the first two minutes until it reached a substantially slower phase of induction, with a maximum NPQ value of ∼1.6 at the end of the first high light treatment. Similarly, during the first transition to darkness, NPQ initially decreased rapidly, followed by a slower phase of relaxation. In the second cycle of high light, NPQ more quickly reached a plateau at a higher level (∼1.7) compared to the last data point of the preceding high light cycle, which could be explained by the accumulation of zeaxanthin in the previous high light phase. The *cp26* plants displayed the same trends but had slightly, though significantly, lower levels than WT plants in the slower phases of NPQ induction (∼95% of WT) and relaxation (∼70% of WT). Chlorophyll fluorescence lifetimes were measured with the same high light/dark protocol but showed no significant differences between the two genotypes. This suggests that some of the NPQ differences observed with PAM fluorescence may have resulted from initial differences in F_m_ rather than differences in the activation of NPQ during light exposure (Fig. 2B).

**Fig. 2:**
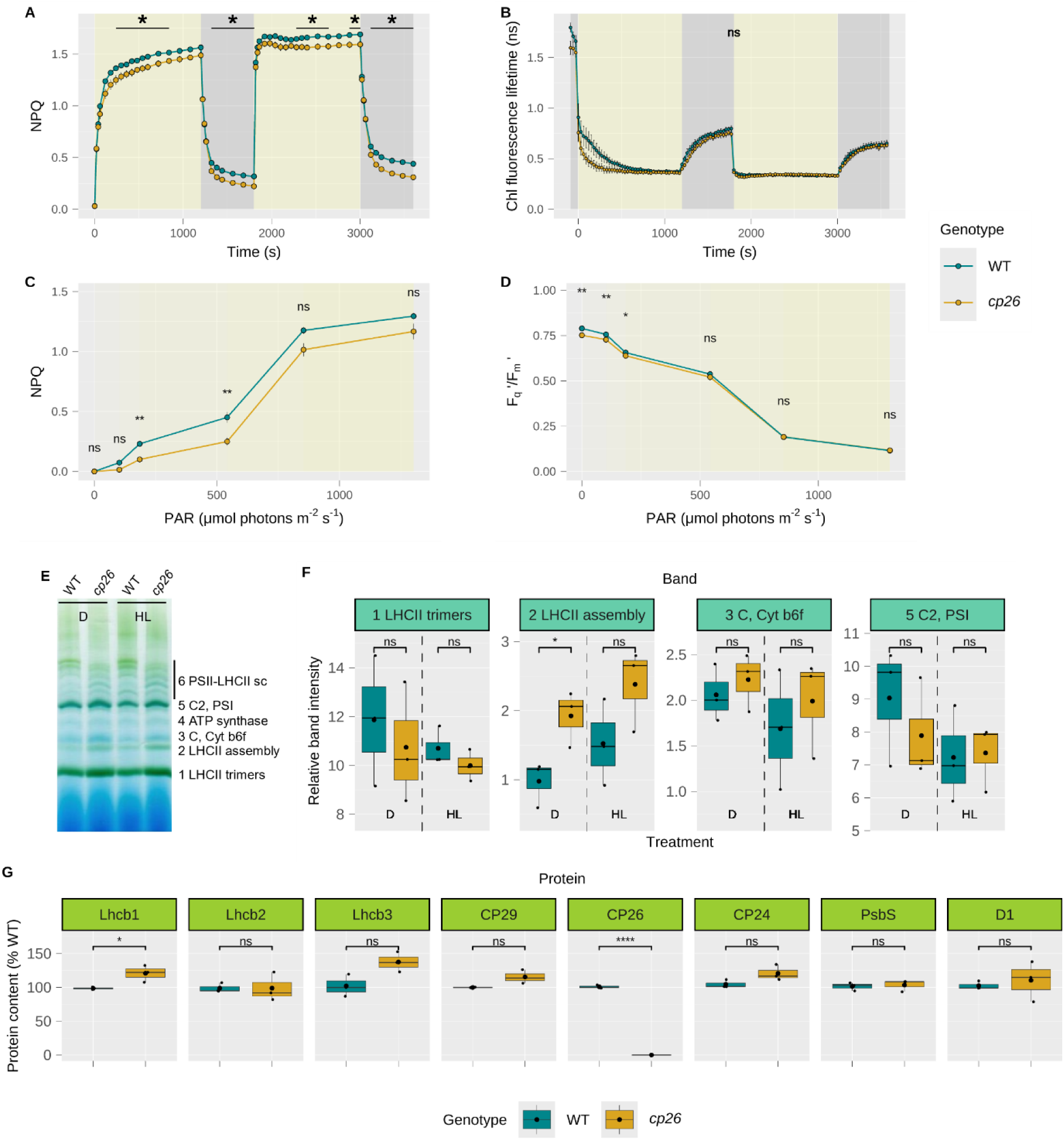
NPQ, PSII quantum yield and PSII antenna composition in *cp26*. **A)** NPQ and **B)** chlorophyll fluorescence lifetime measurements (in nanoseconds) during high light (1000 µmol photons m_-2_ s^-1^ for 20 min; yellow background) and dark phases (10 min; dark grey background) in two consecutive cycles. **C**-**D)** Light response curves from plants exposed to increasing light intensities (photosynthetically active radiation, PAR). PSII parameters (**C** NPQ and **D** PSII efficiency, F_q_’/F_m_’) were calculated by applying a saturating pulse at the end of each light phase. **E**-**G)** Protein complex analyses of thylakoid extracts. **E)** BN-PAGE from dark (D) and high light (HL)-treated samples, representative image from three biological replicates (n=3). **F)** Band intensities corresponding to bands 1-3 and 5 in **E** determined with ImageJ and normalized to the intensity of band 4. **G)** Western blot analyses with ImageJ from three biological replicates (n=3) of high light-treated plants to quantify the abundance of thylakoid membrane proteins relative to the WT. Data in **A**-**D** were collected from six biological replicates (n=6) for PAM fluorescence measurements, and four to six biological replicates (n=4-6) for chlorophyll fluorescence lifetime measurements. Line plots show mean values with error bars indicating the standard error of the mean. The asterisk designates significant differences between WT (green circles) and *cp26* (yellow circles) (two-way repeated measures ANOVA with following paired t-test as post-hoc test). Boxplots (n=3) contain individual data points (black dots), the median (black line inside the box) and the mean (larger black dot inside the box). Significant differences between WT (green boxes/circles) and *cp26* mutant (yellow boxes/circles) are indicated by asterisks (Student’s t-test). Significance levels: ns = no significance, * = p<0.05 and **** = p<0.0001.

To determine at which light level the significant differences in photosynthetic efficiency between WT and *cp26* are lost, chlorophyll fluorescence was also measured in a light response curve (Fig. 2C and 2D). NPQ in *cp26* was generally lower than in the WT upon light exposure, although the difference in NPQ between both genotypes decreased at higher light intensities (Fig. 2C). F_q_’/F_m_’ was significantly lower in *cp26* at low light, but the difference decreased with higher light intensities and both genotypes were similar at light intensities above 200 µmol photons m^-2^ s^-1^ (Fig. 2D).

CP26 is a subunit of the light-harvesting complex that connects strongly bound LHCII trimers to the PSII core. Loss of CP26 has previously been associated with disruption of PSII-LHCII supercomplexes and substitution by other minor antenna proteins, CP29 and CP24 (Andersson *et al*., 2001; de Bianchi *et al*., 2008; Miloslavina *et al*., 2011; Goral *et al*., 2012). To verify these previous observations, thylakoid membrane protein complexes were isolated from dark-acclimated and high light-treated plants, separated in their native state with blue native-polyacrylamide gel electrophoresis (BN-PAGE), and the contents of LHCII proteins determined from high light-treated samples by denaturing Western blot analyses (Fig. 2E-G). The bands on the BN-PAGE gel were associated with protein complexes according to Järvi *et al*. (2011). The largest PSII-LHCII cluster (the top WT band) was missing in *cp26*, whereas the other smaller supercomplexes were more abundant than in the WT, as was band 2 “LHCII assembly”, corresponding to free M-LHCII trimers and the heterodimer CP29/CP24 (Aro *et al*., 2005; Sárvári *et al*., 2022), more so in dark samples than high light samples. Analyses of the relative contents of LHCII proteins in the thylakoid membrane fraction revealed an increase in Lhcb1 (20.8%; p = 0.039), Lhcb3 (37.5%; p = 0.051) and CP29 (15.5%; p = 0.057) protein levels in *cp26* compared to WT (Fig. 2G), although the latter two were not statistically significant.

### *cp26* mutants with varying PsbS levels had lower NPQ than their controls in the slower phase of NPQ induction/relaxation

The potential interaction between CP26 and PsbS was evaluated *in vivo* by crossing the *cp26* mutant with the PsbS knockout line *npq4* (*cp26 npq4*) and the PsbS overexpression line L17 (*cp26 L17*). This allowed comparisons of chlorophyll fluorescence parameters between lines with varying levels of PsbS protein (no PsbS, WT PsbS level and PsbS overexpression) in the presence and absence of CP26, using the same high light/dark protocol described above (Fig. 3).

**Fig. 3:**
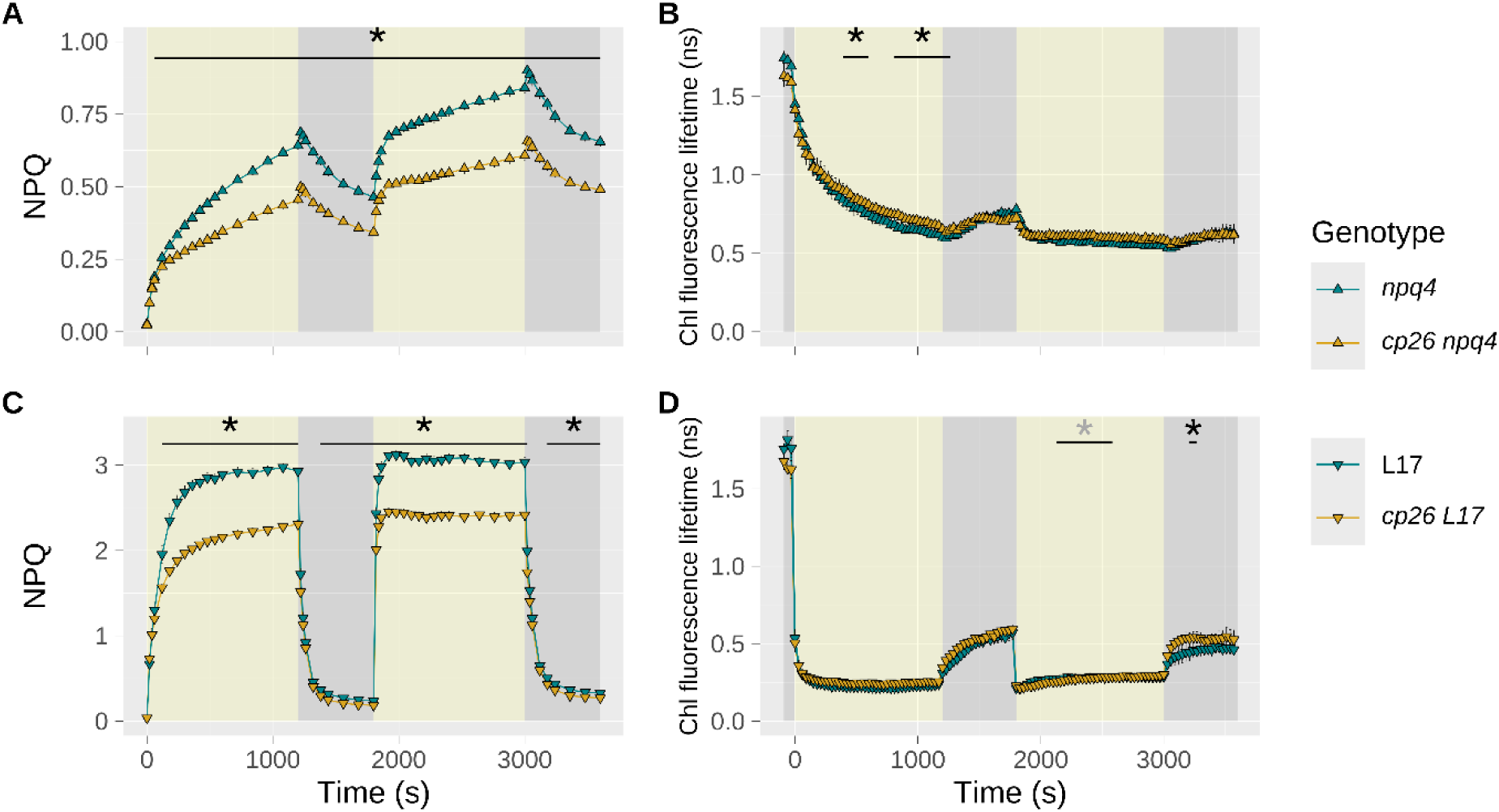
Fluorescence parameters of *cp26* mutants crossed with contrasting PsbS alleles (knockout = *npq4* and overexpression = L17). NPQ (**A** and **C**) and chlorophyll fluorescence lifetime (**B** and **D**, in nanoseconds) measurements of *cp26* double mutants either lacking PsbS (*cp26 npq4*, yellow upwards-pointing triangles in **A** and **B**) or overexpressing PsbS (*cp26 L17*, yellow downwards-pointing triangles in **C** and **D**) and their respective controls *npq4* and L17 (green triangles) during high light induction (1000 µmol photons m^-2^ s^-1^ for 20 min) and dark relaxation (10 min) in two consecutive cycles. Line plots show mean values from six biological replicates (n=6) for PAM fluorescence measurements, and three to six biological replicates (n=3-6) for chlorophyll fluorescence measurements, with error bars indicating the standard error of the mean. The asterisk represents significant differences between two genotypes at each measurement point (two-way repeated measures ANOVA with following paired t-test as post-hoc test). Significance level: * = p<0.05. (The grey asterisk in **D** indicates a region of single significant and non-significant values.)

Consistent with the prominent role of PsbS as a regulator of energy-dependent qE, *npq4* mutants reached approximately 50% lower NPQ than WT during high light induction with minimal relaxation of the established NPQ in the dark (Fig. 3A vs. Fig. 2A). L17 mutants had two-fold higher NPQ amplitudes than WT during high light but reached lower NPQ than WT upon dark relaxation (Fig. 3C vs. Fig. 2A). In both cases, the double mutants *cp26 npq4* and *cp26 L17* displayed significantly lower NPQ than their controls during the slower phase of NPQ induction (after 60 s), which remained lower during both dark relaxation and the following high light/dark cycle in *cp26 npq4*. In PsbS overexpression lines, NPQ levels during the first minute of both dark relaxation phases were not significantly different despite the pronounced difference at the end of the light phase, indicating slower qE relaxation in *cp26 L17*. By the end of the dark phases, however, *cp26 L17* reached lower NPQ values than L17. Curiously, chlorophyll fluorescence lifetime measurements did not show many significantly different values between *cp26* mutants and their controls across the time series (Fig. 3B/D).

Protein complex stoichiometries in the double mutants (Fig. S3) showed similar trends to the analysis of *cp26* (Fig. 2E-G). While most double mutants seemed to have a higher abundance of detached LHCII assemblies relative to their controls (band 2; Fig. S3B), the largest PSII-LHCII supercomplexes visible in *npq4* and L17 were consistently absent upon loss of CP26 (Fig. S3A). Quantitative analyses of the LHCII protein subunits of these samples revealed an increase in Lhcb3 (28.4%; p = 0.092) and CP24 (16.7%; p = 0.018) protein abundance in *cp26 L17* compared to L17 (Fig. S3C) although protein levels varied significantly between the three replicates. Interestingly, the different abundances of PsbS did not seem to affect the stability of different PSII-LHCII supercomplexes.

**Fig. S3:**
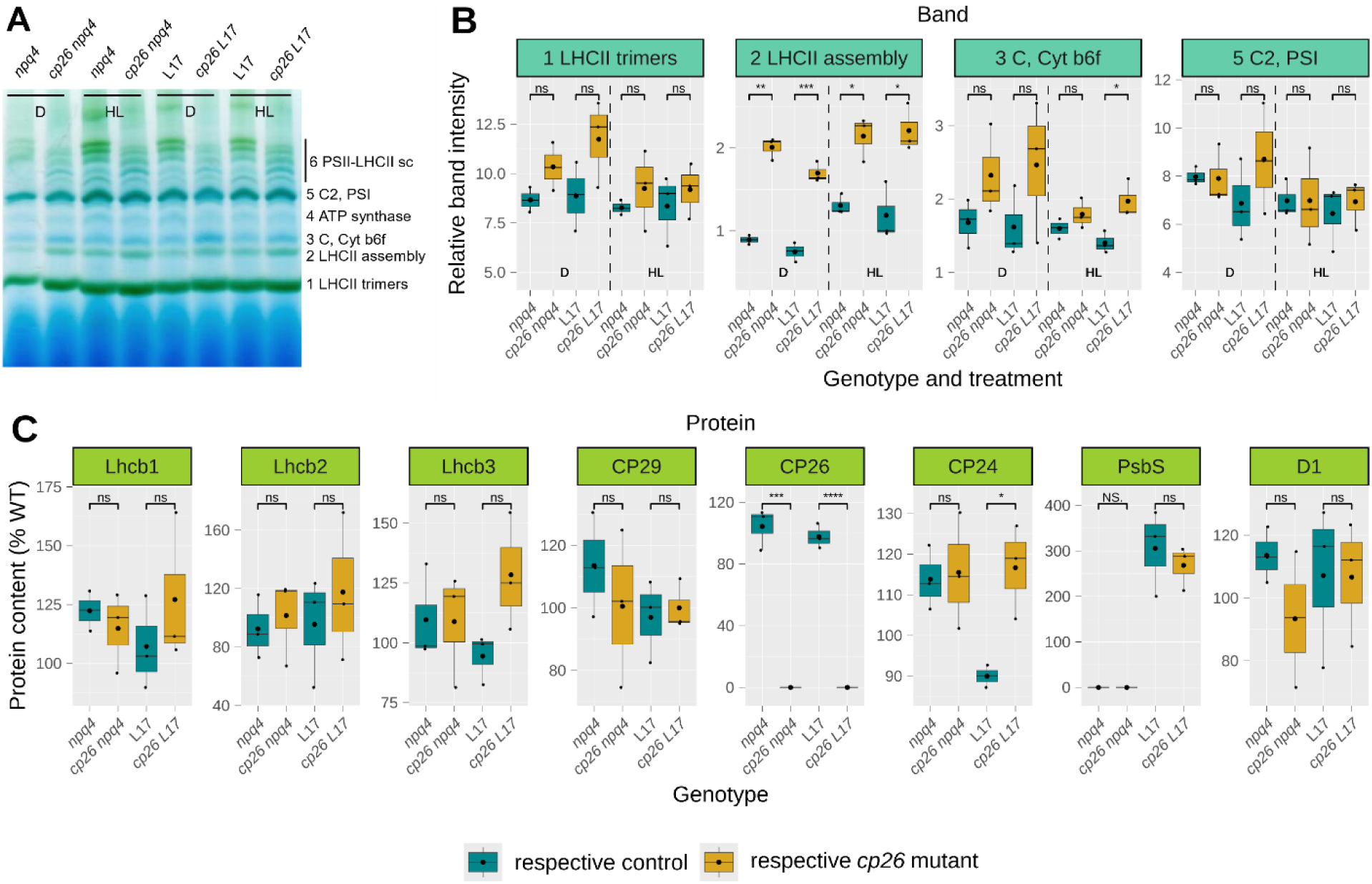
Thylakoid membrane protein analyses of *cp26* mutants crossed with contrasting PsbS alleles (knockout = *npq4* and overexpression = L17). **A)** BN-PAGE of dark (D) and high light (HL)-treated samples from *cp26* double mutants either lacking PsbS (*cp26 npq4*) or overexpressing PsbS (*cp26 L17*) and their respective controls *npq4* and L17; representative image from three biological replicates (n=3). **B)** Band intensities corresponding to bands 1-3 and 5 in **A**, determined with ImageJ and normalized to the intensity of band 4. **C)** Western blot analyses with ImageJ from three biological replicates (n=3) to quantify the abundance of thylakoid membrane proteins relative to the WT (see Fig. 2E). Boxplots (n=3) contain individual data points (black dots), the median (black line inside the box) and the mean (larger black dot inside the box). Significant differences between single mutants (*npq4* and L17, green boxes) and *cp26* mutants (yellow boxes) were determined using Student’s t-test. Significance levels: ns/NS. = no significance and * = p<0.05, ** = p<0.01, *** = p<0.001 and **** = p<0.0001.

### The difference in NPQ due to CP26 deficiency persists when the xanthophyll cycle is blocked

Genetic and pharmacological approaches were used to assess the contribution of the xanthophyll cycle to NPQ in *cp26* (Fig. 4). First, the *cp26* mutant was crossed with the VDE knockout mutant *npq1*, and NPQ was recorded during two cycles of high light/dark phases as described above (Fig. 4A). The *npq1* mutant lacks violaxanthin de-epoxidase (VDE) activity and is, therefore, unable to form zeaxanthin via the reversible violaxanthin cycle. Since zeaxanthin contributes to both qE and qZ components, NPQ levels in this mutant were 50% lower compared to the WT (Fig. 4A vs. Fig. 2A) and had a small decline in the slower phase of NPQ induction. Throughout the high light measurement period, NPQ was significantly lower in the *cp26 npq1* double mutant compared to *npq1,* except during the fast-inducing phases (20-60s), with a more pronounced initial decline in the high light phases (Fig. 4A). Chlorophyll fluorescence lifetime values (Fig. 4B) remained similarly high between high light and dark phases (in contrast to Fig. 2B) and did not differ between genotypes except for the initial dark-acclimated state, again suggesting that the observed differences in NPQ may have arisen in part from quenched F_m_ measurements in plants with the CP26 deficiency.

**Fig. 4:**
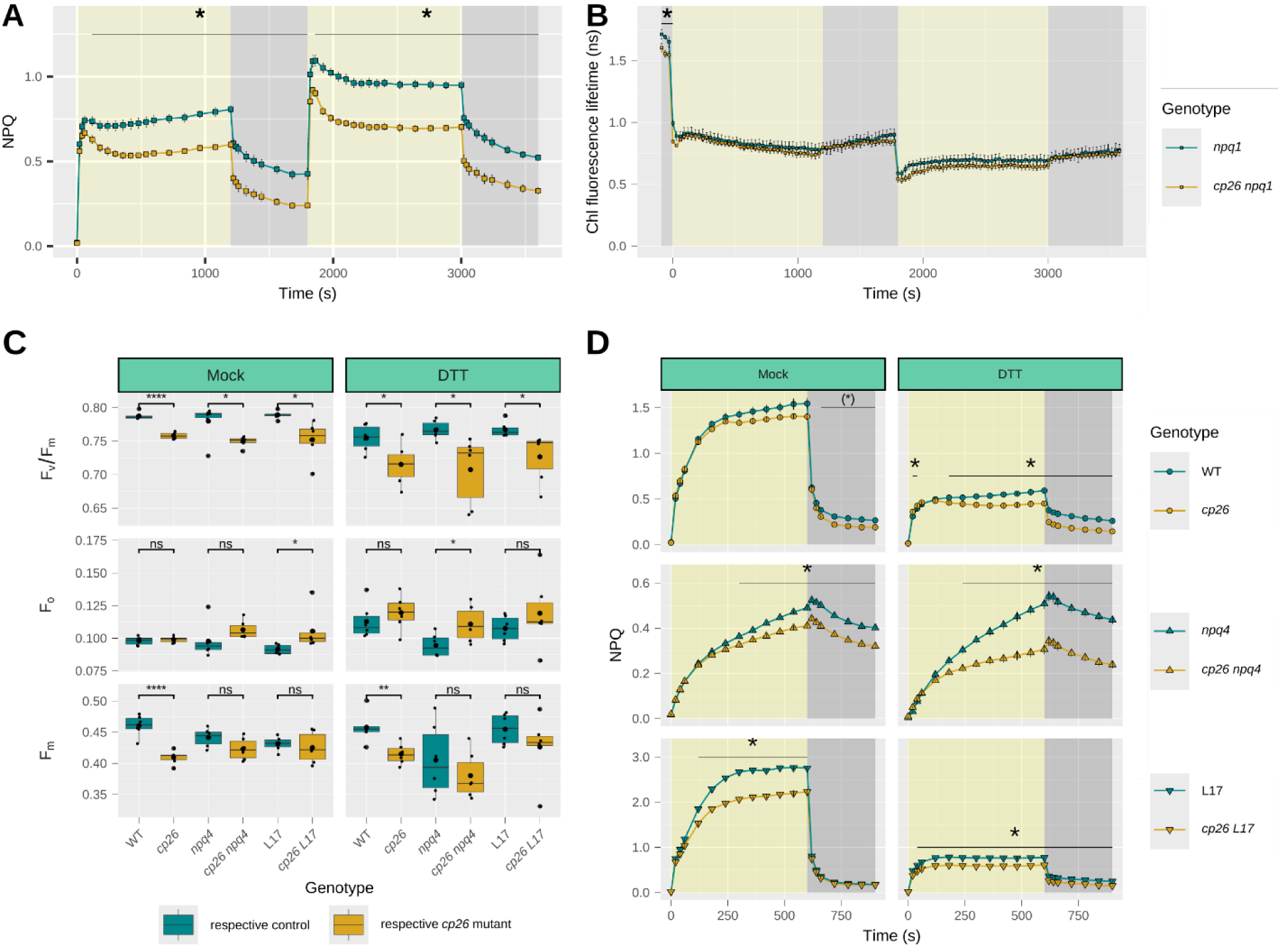
NPQ, chlorophyll fluorescence lifetime and PSII quantum yields in *cp26* mutants and double mutants with contrasting PsbS alleles (knockout = *npq4* and overexpression = L17) infiltrated with DTT or lacking the xanthophyll cycle enzyme VDE (*npq1*). **A)** NPQ and **B)** chlorophyll fluorescence lifetime values (in nanoseconds) of the *cp26* double mutant *cp26 npq1* (yellow squares) compared to the single mutant *npq1* (green squares) during two consecutive high light/dark cycles (20/10/20/10 min). **C-D)** Leaf infiltration assays of WT (green circles), *npq4* (green upward-facing triangles) and L17 (green downward-facing triangles) and respective *cp26* mutants (yellow symbols) using a HEPES/KOH (pH 7.0) buffer as a control (Mock) and the VDE inhibitor DTT. **C)** Maximum quantum yield of PSII, F_v_/F_m_, calculated from minimum (F_o_) and maximum (F_m_) chlorophyll fluorescence levels. **D)** NPQ traces were recorded during one high light (1000 µmol photons m^-2^ s^-1^ for 10 min, yellow background) and one dark cycle (5 min, dark-grey background) by applying saturating pulses. Line graphs show mean values from six biological replicates (n=6) for PAM fluorescence measurements and five to six biological replicates (n=5-6) for chlorophyll fluorescence lifetime measurements, with error bars indicating the standard error of the mean. The asterisk represents significant differences between the two genotypes at each measurement point (two-way repeated measures ANOVA with following paired t-test as post-hoc test). In **D** in plot “Mock/WT vs. *cp26*”, (*) indicates “significant” values although the genotype:time interaction effect was p=0.053. Significance level: * = p<0.05. Boxplots contain individual data points (black dots), the median (black line inside the box) and the mean (larger black dot inside the box). Significant differences between control genotypes (WT, *npq4* and L17; green boxes) and *cp26* mutants (*cp26*, *cp26 npq4* and *cp26 L17*; yellow boxes) are indicated by asterisks (Student’s t-test). Significance levels: ns = no significance, * = p<0.05, ** = p<0.01, *** = p<0.001 and **** = p<0.0001.

Additionally, leaf segments of *cp26*, *cp26 npq4* and *cp26 L17* were infiltrated with dithiothreitol (DTT) diluted in 20 mM HEPES/KOH buffer (pH 7.0) to inhibit VDE activity, and chlorophyll fluorescence was recorded during a shorter 10/5 min high light/dark cycle (Fig. 4C/D). Interestingly, DTT infiltration did not affect the overall trends of the dark values F_v_/F_m_, F_o_ and F_m_ (p = 0.634, 0.215, and 0.065, respectively, two-way ANOVA for genotype:treatment interaction) and NPQ curves compared to the mock treatment (Fig. 4C/D), even though absolute NPQ levels decreased by two-thirds in WT as well as *cp26* and L17 mutants. During the high light phase, the significant decrease in NPQ due to the absence of CP26 appeared to be established somewhat earlier in DTT-treated samples in all three genetic backgrounds (WT, *npq4* and L17), similar to the comparison between *npq1* and *cp26 npq1* (Fig. 4A). These observations clearly showed that the CP26 knockout effect on NPQ persists when the xanthophyll cycle is impaired.

### Inhibition of the proton gradient or blocking of protonatable residues abolished the NPQ induction difference between *cp26* and WT

The above results show that the effect of CP26 knockout on NPQ is independent of the pH sensor protein PsbS. To further investigate the impact of lumen pH on NPQ in *cp26*, the effects of the inhibitors nigericin (Gilmore *et al*., 1995) and *N,N’*- dicyclohexylcarbodiimide (DCCD) (Ruban *et al*., 1992; Li *et al*., 2004) on NPQ in the *cp26* mutants with varying PsbS levels were tested (Fig. 5). Nigericin collapses the proton gradient across the thylakoid membrane, while DCCD binds protonatable protein residues under acidic pH. Measurements of F_v_/F_m_, F_o_ and F_m_ showed that PSII parameters were more strongly affected by DCCD infiltration than nigericin infiltration. Still, the relative changes in F_v_/F_m_, F_o_ and F_m_ due to loss of CP26 trended similarly to the mock infiltration (Fig. 5A) and there was no genotype:treatment interaction effect for any of the three parameters (p = 0.686, 0.772 and 0.298, respectively; two-way ANOVA).

**Fig. 5:**
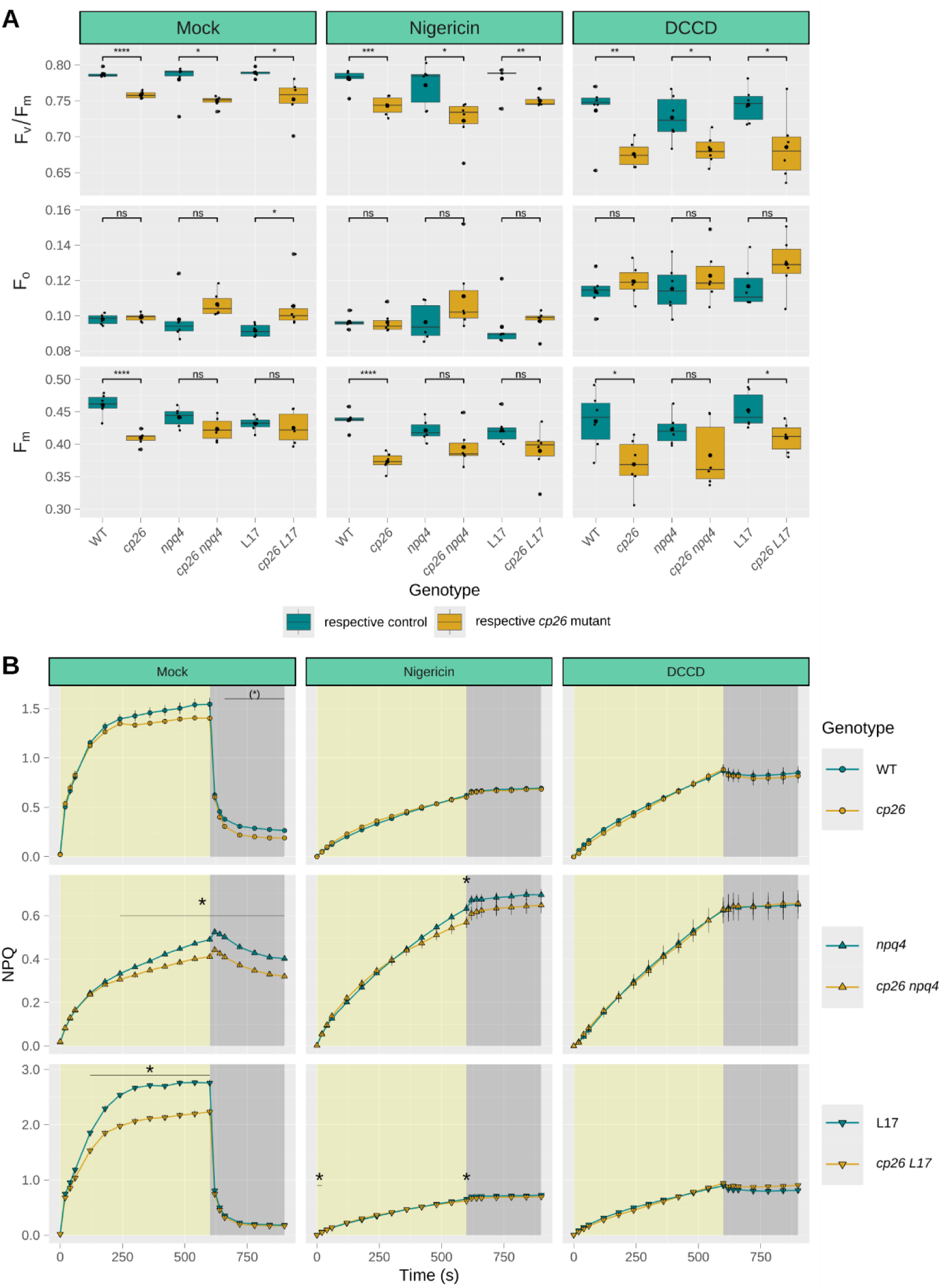
Dark-acclimated chlorophyll fluorescence parameters and NPQ induction in leaves of *cp26* and double mutants with contrasting PsbS alleles (knockout = *npq4* and overexpression = L17) infiltrated with nigericin or DCCD. Leaves were infiltrated with HEPES/KOH (pH 7.0) buffer as a control (Mock, replotted from Fig. 4C/D), nigericin, an inhibitor of the proton gradient across the thylakoid membrane, and DCCD, an inhibitor of protonatable protein residues in the thylakoid lumen. **A)** Maximum quantum yield of PSII, F_v_/F_m_, calculated from minimum (F_o_) and maximum (F_m_) chlorophyll fluorescence levels. **B)** NPQ was recorded for the genotype pairs: WT vs. *cp26* (green and yellow circles, respectively); *npq4* vs. *cp26 npq4* (green and yellow upwards-pointing triangles, respectively); and L17 vs. *cp26 L17* (green and yellow downwards-pointing triangles, respectively) during one high light (1000 µmol photons m^-2^ s^-1^ for 10 min, yellow background) and dark cycle (5 min, dark-grey background). Boxplots contain individual data points (black dots), the median (black line inside the box) and the mean (larger black dot inside the box) from six biological replicates (n=6). Significant differences between control genotypes (WT, *npq4* and L17; green boxes) and *cp26* mutants (*cp26*, *cp26 npq4* and *cp26 L17*; yellow boxes) are indicated by asterisks (Student’s t-test). Significance levels: ns = no significance, * = p<0.05, ** = p<0.01, *** = p<0.001 and **** = p<0.0001. Line graphs show mean values from six biological replicates (n=6) with error bars indicating the standard error of the mean. The asterisk represents significant differences between the two lines at each measurement point (two-way repeated measures ANOVA with following paired t-test as post-hoc test when applicable). In **B** in plot “Mock/WT vs. *cp26*”, (*) indicates “significant” values although the genotype:time interaction effect was p=0.053. Significance levels: * = p<0.05.

Acidification of the thylakoid lumen is a prerequisite for the qE component of NPQ, transduced through protonation of glutamate residues in PsbS. Collapsing this pH gradient using nigericin, should therefore mimic a PsbS knockout. Indeed, NPQ traces of all genotypes treated with nigericin were highly comparable to the *npq4* mock infiltration (Fig. 5B), reaching NPQ levels of ∼0.7 at the end of the high light phase and showing negligible NPQ relaxation in the dark. Interestingly, nigericin and DCCD infiltrations abolished the differences in NPQ between WT CP26 and *cp26* alleles, despite the pronounced impact of CP26 deficiency on the dark-acclimated fluorescence parameters. This suggests that the reduction in NPQ observed in *cp26* mutants with PAM-fluorescence may not originate entirely from a partially pre-quenched F_m_ in dark-acclimated samples - acidification of the thylakoid lumen may also play a role, perhaps via protonation of lumen-exposed protein residues.

### Proton motive force estimates are not affected in *cp26* mutants

To find out if differences in NPQ induction between single and *cp26* double mutants arose due to an effect of the CP26 deficiency on the formation of the trans-thylakoid ΔpH, the proton motive force was assessed *in vivo* via electrochromic shift measurements. However, no clear effects on the proton conductivity of the ATP synthase, gH^+^ (Fig. 6A), the steady-state proton flux rate, vH^+^ (Fig. 6B) or the overall proton motive force (pmf, Fig. 6C) were observed for the single *cp26* mutant or the three double mutants (*cp26 npq4*, *cp26 L17* and *cp26 npq1*).

**Fig. 6:**
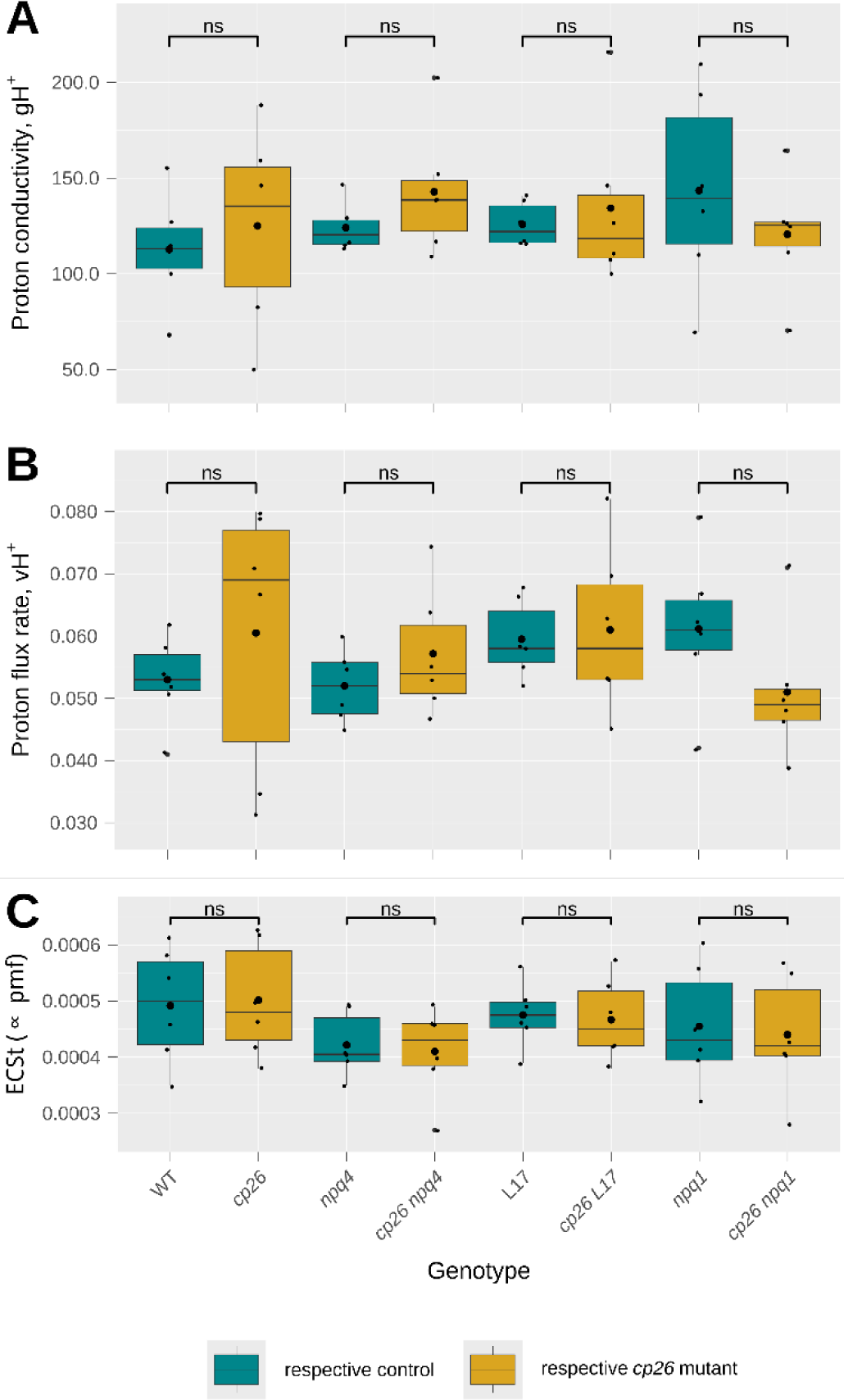
Evaluation of the proton gradient across the thylakoid membrane. **A)** ATP synthase proton conductivity, gH^+^, **B)** steady-state proton flux rate, vH^+^, and **C)** ECSt (electrochromic shift signal during light-to-dark transition; proportional to trans-thylakoid proton motive force, pmf, Baker *et al*., 2007) were measured from six biological replicates (n=6) using the MultispeQ (Photosynq) and a modified RIDES protocol with a light intensity of 1000 µmol photons m^-2^ s^-1^. Boxplots contain individual data points (black dots), the median (black line inside the box) and the mean (larger black dot inside the box). Significant differences between *cp26* mutants (*cp26*, *cp26 npq4*, *cp26 L17* and *cp26 npq1*; yellow boxes) and their respective controls (WT, *npq4*, L17 and *npq1*; green boxes) were determined using Student’s t-test (significance levels: ns = no significance).

## Discussion

The minor LHCII antennae CP29, CP26 and CP24 connect the major LHCII trimers to the PSII core and thereby facilitate excitation energy transfer to the reaction centers, whilst also contributing to light harvesting and NPQ formation (Jansson, 1999). Of these three proteins, CP29 has been the main target of NPQ research (Feng *et al*., 2013; Betterle *et al*., 2015, 2017; Gacek *et al*., 2020; Guardini *et al*., 2020, 2022; Mascoli *et al*., 2020; Cignoni *et al*., 2021; Elias *et al*., 2023; Sardar *et al*., 2022, 2024) as its deletion showed an initial impact on NPQ induction upon high light exposure (de Bianchi *et al*., 2011; Miloslavina *et al*., 2011; Guardini *et al*., 2020). The CP24 protein is highly dependent on the presence of CP29, and a *cp24* knockout mutant was also impaired in overall NPQ capacity (Kovács *et al*., 2006; de Bianchi *et al*., 2008; Chen *et al*., 2018). On the other hand, the deletion of CP26 was found to have only a minor influence (de Bianchi *et al*., 2008), although *in vitro* studies of CP26 demonstrated a unique connection to and dependency on the xanthophyll zeaxanthin (Morosinotto *et al*., 2002; Dall’Osto *et al*., 2005).

### The quenched dark state in *cp26* mutants is likely due to a less stable PSII-LHCII supercomplex architecture

Analysis of the maximum PSII quantum efficiency after dark acclimation revealed that *cp26* mutants had lower F_v_/F_m_ values compared to their controls (Fig. 1A). This was mainly due to shorter chlorophyll fluorescence lifetimes (Fig. 1D) causing a decrease in F_m_ (Fig. 1C), and only a slight increase in F_o_ (Fig. 1B), suggesting that these mutants were already in a dissipative state before the onset of light. These observations are consistent with previous observations for mutants lacking minor antenna proteins (de Bianchi *et al*., 2008, 2011; Chen *et al*., 2018), and indicate a structural impairment rather than specific roles of these proteins. Under dark conditions, the PSII core and LHCII proteins form large supercomplexes to maximize light interception when photons are limited. By deleting CP26, the strongly bound S-LHCII trimers are no longer connected to the PSII core and potentially form aggregates in the thylakoid membrane, which could explain the pre-quenched state in *cp26* (Horton *et al*., 1991; Mullineaux *et al*., 1993; Miloslavina *et al*., 2008, 2011; Goral *et al*., 2012; Tian *et al*. 2015; Natali *et al*., 2016; Adams *et al*., 2018; Shukla *et al*., 2020; Tutkus *et al*., 2021; Wilson *et al*., 2022). BN-PAGE analysis of *cp26* (Fig. 2E/F and Fig. S2A/B) clearly showed the absence of the highest order PSII-LHCII supercomplexes and a trend towards more abundant free LHCII assemblies (M-LHCII/CP29/CP24; Aro *et al*., 2005; de Bianchi *et al*., 2011; Sárvári *et al*., 2022) upon solubilization, which is consistent with previous observations of less stable supercomplexes in CP26 antisense lines (Yakushevska *et al*., 2003), as well as *cp29* and *cp24* knockout mutants (de Bianchi *et al*., 2008, 2011; Miloslavina *et al*., 2011; Dall’Osto *et al*., 2017; Cazzaniga *et al*., 2020; Ilíková *et al*., 2021; Guardini *et al*., 2022). If LHCII aggregation is the reason for the dark-quenched state in *cp26*, it should be independent of the prerequisites of energy-dependent quenching qE, i.e. lumen acidification and zeaxanthin formation. Indeed, when these were inhibited, either genetically in the double mutants *cp26 npq4* and *cp26 npq1*, or via leaf infiltration with nigericin or DCCD (Fig. 4 and Fig. 5), the dark-quenched state in *cp26* persisted.

Upon high light exposure, the antenna architecture is thought to be rearranged by the action of the PsbS protein and the M-LHCII/CP29/CP24 subcomplex is disconnected from the PSII core, thereby decreasing the antenna size around PSII alongside the activation of NPQ (Bergantino *et al*., 2003; Kiss *et al*., 2008; Wilk *et al*., 2013; Sacharz *et al*., 2017). Surprisingly, BN-PAGE analysis did not display any differences in PSII-LHCII supercomplex assemblies between dark and high light samples (p = 0.416, one-way ANOVA) or *npq4* and L17 samples (p = 0.981, one-way ANOVA) (Fig. 2E and Fig. S3A). This seems to be a common issue in some of the literature (Suorsa *et al*., 2015 and Pashayeva *et al*., 2021 but not Dong *et al*., 2015) and shows the limitations of using detergents for membrane solubilization, disrupting weak protein interactions more easily. Nevertheless, PSII operating efficiency over a range of increasing light intensities showed that *cp26* only had lower F_q_’/F_m_’ values under low light conditions up to about 200 µmol photons m^-2^ s^-1^ (Fig. 2D). These results indicate that the loss of energy transfer efficiency due to the absence of CP26 may be less important under high light conditions where antenna size is generally downregulated to avoid overexcitation of PSII.

Previous research on the *cp26* SALK line used here (de Bianchi *et al*., 2008) suggested that the lack of CP26 might be complemented by upregulation of other minor antenna proteins. We did see a similar non-significant increase in CP29 in addition to Lhcb1 and Lhcb3 in *cp26* and a slight increase in CP24 in the *cp26 L17* double mutant (Fig. 2G and S3C), all of which are subunits of the M-LHCII/CP29/CP24 assembly, corroborating the BN-PAGE data with a more abundant band 2 on the gels (Fig. 2 and S3). However, these results are not consistent between the different genotype pairs and the Western Blot analyses varied substantially between replicates. This, together with the clear lack of higher order supercomplexes in *cp26* genotypes in BN-PAGE results, demonstrates that despite the functional and conformational similarity between different LHCII proteins (Croce and van Amerongen, 2011), deficiency of CP26 is not complemented by the function of another LHCII protein.

### CP26 contributes to NPQ in the slower phases of NPQ induction/relaxation

NPQ is typically calculated according to the Stern-Volmer equation NPQ = (F_m_/F_m_’) – 1 and is therefore dependent on the maximum chlorophyll fluorescence value F_m_, which is initially measured after dark acclimation. As discussed above, F_m_ is significantly lower in the *cp26* mutant, suggesting that some form of NPQ is already active in the dark, and could therefore be a confounding factor for NPQ data in the light. A theoretical model that assumes a standard F_v_/F_m_ value of 0.83 and calculates NPQ(T) (Tietz *et al*., 2017), consistently shifts NPQ values of the *cp26* mutants to higher levels than their controls (except for *cp26 L17*, data not shown). However, this makes it difficult to distinguish between dark and different light-induced NPQ events. Thus, it is important to consider that the dark-quenched state in the *cp26* mutant is reflected in F_m_ and the relationship between F_m_ and F_m_’ only reflects further NPQ induction in response to light. Therefore, it is interesting that the initial rise in NPQ upon high light induction fully overlapped between *cp26* and WT and NPQ was only lower in the former after a minute of high light exposure. The same trend also applied to the dark relaxation phase and the second cycle, despite the previously accumulated differences in NPQ. The first minute of NPQ induction is typically attributed to the qE component, followed by the slowly induced qZ and qI components (Nilkens *et al*., 2010). This could suggest that qE is not affected by the absence of CP26, but that CP26 may contribute to one of the latter two mechanisms (or an alternative unexplored NPQ mechanism). As CP26 protein conformation was shown to be dependent on the xanthophyll zeaxanthin (Morosinotto *et al*., 2002; Dall’Osto *et al*., 2005), it could be speculated that CP26 plays a role in qZ (Miloslavina *et al*., 2011).

Significant differences in NPQ between the *cp26* mutant and WT were mainly observed during dark relaxation phases, which conforms with previously published studies (de Bianchi *et al*., 2008). These differences seemed to increase in amplitude with the subsequent light/dark cycles (Fig. 2A). Nevertheless, chlorophyll fluorescence lifetime measurements, which reflect all NPQ processes combined, showed no significant differences between the two genotypes (Fig. 2B), possibly also due to different measuring light wavelengths being employed for NPQ (540 nm) and lifetime (∼404 nm) measurements. These results indicate that the overall decrease in NPQ induction in *cp26* in response to light is likely dependent on the pre-quenched F_m_.

### Absence of PsbS and VDE exacerbates NPQ induction differences in *cp26* mutants

Experiments with *cp26* double mutants with differing levels of PsbS (Fig. 3A/C) and VDE (Fig. 4A) as well as with leaf infiltration with the VDE inhibitor DTT (Fig. 4D) showed that the differences in NPQ during the slower phases of induction/relaxation between *cp26* mutants and their respective controls persisted and were extended to the fast phases of relaxation despite the absence of energy-dependent qE and zeaxanthin-dependent qZ. These results indicate that PsbS and VDE do not act upstream of the CP26 effect on NPQ induction (otherwise NPQ traces would have converged throughout the measurements in the absence of either protein). Notably, the NPQ differences are more pronounced upon the alteration of qE components in contrast to the WT vs. *cp26* comparison (Fig. 2A). This may suggest that this NPQ component becomes more visible under strong lumen acidification when the moderating effect of qE on lumen pH is removed. These findings are partially in line with *in vitro* results that demonstrated PsbS-independent conformational changes in CP26 (Dall’Osto *et al*., 2005). However, in the results by Dall’Osto *et al*. (2005), these conformational changes required the presence of zeaxanthin, which we could not confirm here.

### The impact of CP26 on NPQ induction requires the proton gradient and depends on the protonation of lumen-exposed residues

Thylakoid lumen acidification through build-up of a proton gradient is an important activator of NPQ. Evaluation of the steady state proton gradient in *cp26* mutants (Fig. 6) did not show an impairment compared to the controls, which aligns with previous observations using 9-aminoacridine (de Bianchi *et al*., 2008). Consistent with the role of thylakoid lumen acidification on NPQ induction, leaves infiltrated with nigericin, which collapses the proton gradient across the thylakoid membrane, no longer showed any difference in NPQ induction between *cp26* mutants and their controls. This was further corroborated by DCCD infiltration, which blocks protonatable protein residues in the thylakoid lumen (Ruban *et al*., 1992; Walters *et al*., 1996), again abolishing the contribution of the CP26 mutation on NPQ induction in all single and double mutants (Fig. 5B). Together, these findings suggest that the NPQ capacity of CP26 depends on lumen acidification, potentially via protonation of specific residues on CP26. Interestingly, previous *in vitro* studies detected two glutamate residues in CP26 that were blocked by binding DCCD (Walters *et al*., 1996), but further *in vivo* analyses are required to test a putative role in NPQ induction. Targeted point mutational studies of similar protonatable residues in CP29, however, did not impact CP29-specific NPQ effects and were deemed irrelevant to NPQ induction (Guardini *et al*., 2020).

## Conclusions

The involvement of the minor antennae in NPQ is still debated (Xu *et al*., 2015; Dall’Osto *et al*., 2017; Townsend *et al*., 2018; Saccon *et al*., 2020; Ruban and Wilson, 2021). There is evidence that a small proportion of overall NPQ may be associated with minor antennae quenching (Miloslavina *et al*., 2011; Dall’Osto *et al*., 2017), while the main quenching site has been proposed to be in detached major LHCII trimers (Nicol *et al*., 2019). In algae and mosses, however, it was recently reported that CP26 is highly important for NPQ induction in the macroalga *Ulva* (Gao *et al*., 2020), the green alga *Chlamydomonas* (Cazzaniga *et al*., 2020, 2023) and the moss *Physcomitrella* (Peng *et al*., 2019), with contributions up to 70% in NPQ induction. NPQ in chlorophyte algae and mosses, however, differs from NPQ mechanisms in higher plants and depends on light-harvesting complex stress-related (LHCSR) proteins (Peers *et al*., 2009). During the evolution of higher plants, PsbS became the main player of qE induction, which possibly changed NPQ dynamics of the minor antennae.

Here we followed up on some long-standing hypotheses regarding the role of CP26 in NPQ. We confirmed that a CP26 knockout mutation leads to a dark-quenched state (Nicol *et al*., 2019) and less stable antennae, which impair photochemical efficiency in darkness and low light below ∼200 µmol photons m^-2^ s^-1^. We also observed lower NPQ induction in *cp26* and confirmed that this effect is not dependent on qE components PsbS, as postulated by Dall’Osto *et al*. (2005), and zeaxanthin, despite the *in vitro* evidence that conformational changes in CP26 are dependent on the xanthophyll zeaxanthin (Morosinotto *et al*., 2002; Dall’Osto *et al*., 2005). Instead, it might involve protonation of lumen-exposed glutamate residues on CP26 (Walters *et al*., 1996) upon lumen acidification, potentially leading to the conformational change that switches CP26 into a quenching state with lutein possibly as the quencher (Avenson *et al*., 2009). Complementation of the *cp26* mutant with CP26 sequences mutated in putative NPQ sites will help to disentangle the dual function of CP26 as a structural PSII-LHCII supercomplex component and a possible NPQ component and contribute to our understanding of the intricate NPQ network.

## Methods

### Plant material and growth conditions

All mutant lines were generated from the *Arabidopsis* wild-type Columbia-0 (Col-0). *Arabidopsis* mutant lines *cp26* (SALK-014869C, N656198), *npq4* (*PsbS*, N66021) and *npq1* (*VDE*, N3771) were obtained from the Nottingham Arabidopsis Stock Centre (NASC). The PsbS overexpression line L17 was generated by the Niyogi lab (Li *et al*., 2002). Double mutants *cp26 npq4*, *cp26 L17* and *cp26 npq1* were generated by cross-pollination and homozygous mutant lines were selected in the F2 generation by chlorophyll fluorescence screen under high light in the FluorCam imager (Photon Systems Instruments, Czech Republic) and by PCR screen using the primers LBb1.3/AT cp26_LP/AT cp26_RP (*cp26*), KN118/KN119 (*npq4*, Li *et al*., 2000), AT PsbS_2_S/AT scpl16_fw/AT scpl16_rv (L17), and KN75/KN76 (*npq1*, Niyogi *et al*., 1998) (Table S1). All mutants were confirmed on DNA and protein levels (Fig. S1). The primers for L17 genotyping were developed by mapping the T-DNA insertion site for L17 to exon 11 of the serine carboxypeptidase-like 16 gene (*SCPL16*, AT3G12220) using TAIL-PCR (Singer and Burke, 2003; see Fig. S4). All plants were initially grown for 4-6 weeks under short-day-conditions (8-h light/16-h dark, 22°C, 60% humidity, 150 µmol m^-2^ s^-1^) and then shifted to a growth cabinet with a 12-h light/dark cycle and similar conditions at least two days before experiments.

**Fig. S4:**
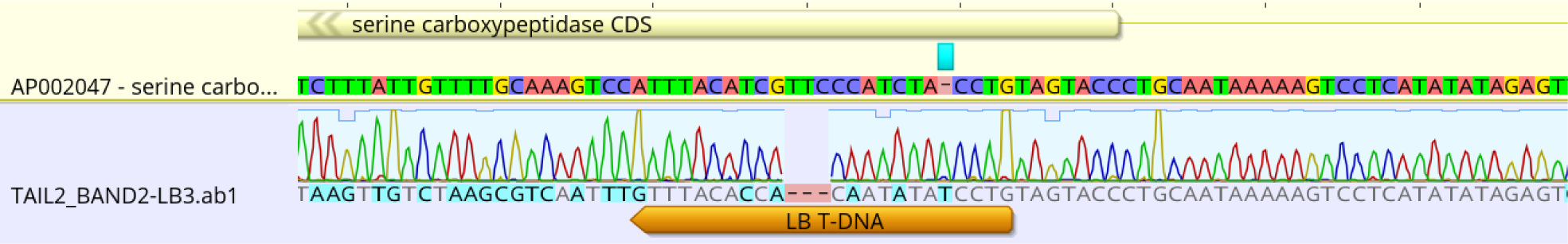
L17 T-DNA insertion site. The sequenced PCR product resulting from TAIL-PCR was mapped to the coding sequence of exon 11 of the serine carboxypeptidase-like 16 gene (*SCPL16*, AT3G12220) using Geneious Prime. The left border T-DNA repeat sequence in the TAIL-PCR product (LB T-DNA) and the putative T-DNA insertion site in the *SCPL16* gene (light turquoise rectangle) are highlighted.

### Chlorophyll fluorescence measurements

Chlorophyll fluorescence was measured using the Dual-Klas-NIR instrument (Walz, Germany) coupled to a GFS-3000 measuring chamber and a LI-6800 gas exchange system (LI-COR, USA). Temperature was controlled at 25°C via the measuring chamber and CO_2_ (410 ppm) and humidity (60%) were controlled via the LI-6800 console. Whole plants were dark-acclimated for 75 min. Leaves clamped in the cuvette were exposed to two cycles of 20 min high light (1000 µmol photons m^-2^ s^-1^ red actinic light) and 10 min dark relaxation. Chlorophyll fluorescence was measured using weak green PAM measuring light (540 nm). PSII efficiency parameters were determined by applying a saturating pulse (4000 µmol photons m^-2^ s^-1^ red actinic light, 800 ms) directly after dark acclimation to measure the maximum PSII efficiency (F_v_/F_m_=(F_m_-F_o_)/F_m_), followed by a 10-s dark period before switching on actinic light, and by applying further saturating pulses during high light exposure and dark relaxation to calculate non-photochemical quenching (NPQ=(F_m_-F_m_’)/F_m_’) and PSII operating efficiencies (F_q_’/F_m_’=(F_m_’-F’)/F_m_’).

For infiltration with inhibitors, leaf segments of 1.5 cm x 1.5 cm were dark-acclimated on wet filter paper for 60 min, then vacuum-infiltrated in a syringe and briefly dried on filter paper (under dark conditions) and further dark-acclimated for 15 min inside the GFS-3000 measuring chamber before running a protocol of 10 min high light and 5 min dark relaxation with the same settings as described above. All inhibitors were dissolved in HEPES buffer (20 mM HEPES/KOH, pH 7.0). For removal of the proton gradient across the thylakoid membrane, leaves were infiltrated with 50 µM nigericin sodium salt (Sigma Aldrich; dissolved in ethanol). For blocking protonatable protein residues, leaves were infiltrated with 30 mM *N,N′*-dicyclohexylcarbodiimide (DCCD, Sigma Aldrich; dissolved in dimethylformamide, DMF). The VDE protein was inhibited by infiltration with 5 mM dithiothreitol (DTT, Sigma Aldrich). As a control, leaves were infiltrated with buffer only (mock infiltration).

Light curves were measured on intact leaves after 75 min dark acclimation. Light intensities were sequentially increased from low to high (0, 102, 185, 542, 853, 1302 µmol photons m^-2^ s^-1^) in steps of 5 min with a saturating pulse at the end of each step to calculate chlorophyll fluorescence parameters.

### Fluorescence lifetime snapshot measurements

Time-correlated single photon counting (TCSPC) was used to measure the initial chlorophyll fluorescence lifetimes and the changes in response to the high light and dark periods, as previously described (Steen *et al*., 2020). A Ti:sapphire oscillator (Coherent, Mira900f, 76 MHz) generated pulses at ∼808 nm and were frequency-doubled to ∼404 nm, which was used to excite the Soret band of chlorophyll *a*. This excitation beam was then divided by a beamsplitter, part of which was directed into a photodiode (Becker-Hickl, PHD-400) to provide SYNC signals. The other half of the excitation beam was then incident at an approximately 70° angle to the adaxial side of the leaf lamina. The excitation power was set to 1.0 mW, with a ∼600 µm beam diameter, which is enough to saturate the reaction centers. The samples were exposed to an actinic light (Leica KL1500 LCD) sequence, composed of alternating high light (1000 µmol photons m^-2^ s^-1^) and dark periods of 20-10-20-10 min. During this illumination sequence, snapshots of chlorophyll fluorescence lifetime were taken every 30 s. Fluorescence photons were detected by a microchannel plate (MCP)-photomultiplier tube (PMT) detector (Hamamatsu R3809U MCP-PMT) following a monochromator (HORIBA Jobin-Yvon; H-20), which was set to 680 nm, specifically detecting chlorophyll *a* Qy band fluorescence. A LabVIEW program was used to control a series of shutters, thereby coordinating the application of the excitation beam, the actinic light, and the detector. Within a 1-s total integration time of detection, a 0.2-s portion of the data showing the longest lifetime was selected for further data processing to ensure the saturation of the reaction centers (Sylak-Glassman *et al*., 2016). Each fluorescence decay profile was fitted with a bi-exponential decay function and the amplitude-weighted average lifetime was calculated by:

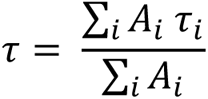

where *A*_*i*_ and *τ*_*i*_ are the amplitudes and fluorescence lifetimes of the i^th^ fitting component, respectively.

### Electrochromic shift measurements

Steady state trans-thylakoid proton conductivity and proton motive force were estimated *in situ* on light-acclimated plants inside the growth cabinet with a hand-held MultispeQ device (Photosynq) using the protocol “Photosynthesis RIDES 2.0 EC 1000-no-open” (gH^+^, vH^+^ and ECSt parameters).

### Blue-native polyacrylamide gel electrophoresis (BN-PAGE) and Western blotting

Thylakoid membrane proteins were isolated from 8-week-old plants according to Järvi *et al*. (2011). One plant per genotype was either treated with 1000 µmol photons m^-2^ s^-1^ white high light or dark-acclimated for one hour, with three biological replicates each.

Fresh whole rosettes were ground in 20 ml ice-cold grinding buffer either under high light or in darkness and filtered through two layers of Miracloth (Sigma Aldrich). Extracts were further treated on ice and in dim light by several centrifugation steps and resuspension of pellets in shock and storage buffers. All buffers contained 10 mM NaF to preserve protein phosphorylations. After the final resuspension step in 200 µl of storage buffer, the chlorophyll *a* content was determined in 100% methanol according to Porra *et al*. (1989), using absorption values at wavelengths 652.0 nm, 665.2 nm and 750 nm for the equation: chlorophyll *a* [μg/ml] = 16.29 x (A_665.2_ - A_750_) – 8.54 x (A_652.0_ - A_750_).

For BN-PAGE analysis, extracts were resuspended with freshly prepared ice-cold ACA buffer (containing 10 mM NaF) to a concentration of 1 mg chlorophyll/ml. Aliquots of 5 µg chlorophyll were resuspended with an equal volume of 4% digitonin (Sigma Aldrich) in ACA buffer (final concentration of 2% digitonin), and thylakoid membrane proteins were solubilized at room temperature for 10 min on a shaker. Afterwards, samples were spun down for 20 min at 18 000*xg* at 4°C to pellet insolubilized membranes. The supernatant was mixed carefully with 1/10 of the volume with sample buffer, loaded onto a 3-12% Bis-Tris NativePAGE gradient gel (Invitrogen) and the electrophoresis was run on ice for five hours with increasing voltage (75 – 150 V).

Protein abundance was compared between genotypes using thylakoid samples in Western blot analyses. Samples equivalent to 1 µg chlorophyll were solubilized with 2X sample buffer (Laemmli buffer + 6M urea + β-mercaptoethanol + Bromophenol blue) for 5 min at 75°C and loaded onto 12% Mini-PROTEAN® TGX™ Precast Gels (Bio-Rad). After protein separation, proteins were blotted onto PVDF membranes with a Trans-Blot Turbo Transfer System (Bio-Rad). Blots were washed in TBS, blocked with 5% milk in T-TBS for one hour, washed twice with T-TBS and incubated overnight with primary antibodies in 1% milk in T-TBS shaking at 4°C (dilutions according to manufacturer’s instructions). The day after, blots were washed four times with T-TBS, incubated in secondary antibody (goat anti-rabbit IgG, HRP conjugated, dilution = 1:12500 in 1% milk in T-TBS) for one hour and washed twice with T-TBS and three times with TBS. For the detection of protein bands, blots were incubated in ECL solution for 5 min, and imaged with a G:Box Chemi XRQ system (Syngene). Image analysis and quantification of bands were performed in ImageJ, using a dilution series of WT samples as a standard on each blot. All primary and secondary antibodies were obtained from Agrisera/Newmarket Scientific (Sweden/UK).

### Statistical analyses

All statistical analyses were performed in R (version 4.4.0) using the packages “rstatix”, “ggpubr” and “car”. Time course datasets were first tested for extreme outliers and normal distribution of data points for each genotype at each time point (Shapiro-Wilk test). A two-way repeated measures ANOVA was conducted to determine the effects of genotype and time on NPQ, F_q_’/F_m_’, F_v_’/F_m_’ and chlorophyll fluorescence lifetime data. If a significant interaction effect (genotype:time) was detected, a paired t-test with Bonferroni-adjusted p-values was performed to obtain the genotype effect between *cp26* mutants and their respective controls at each time point (Supplemental data file).

For datasets presented in boxplots, a Student’s t-test was used to compare the means of two genotypes (*cp26* mutants vs. their controls) using the ggplot2 package. Before that, Levene’s test was used to check for equality of group variances.

## Acknowledgements

The authors thank Dr. Rich Vath and Dr. Cris Sales for help with the integration of the GFS-3000 cuvette with the LI-6800 console. For the purpose of open access, the authors have applied a Creative Commons Attribution (CC BY) license to any Author Accepted Manuscript version arising from this submission.

## Author contributions

JW and JK conceived the study, JW and JK designed the experiments, JW carried out all experiments, data analysis and interpretation, except for TCSPC measurements, LL conducted TCSPC measurements with assistance from DP-T and AM. GT helped map the L17 T-DNA insertion site. All authors contributed to the manuscript and approved the submitted version.

## Supplemental data

Table S1: Primers used in this study.

Fig. S1: Molecular confirmation of *Arabidopsis* mutants used in this study.

Fig. S2: PSII quantum efficiencies in *cp26* and double mutants with contrasting PsbS (knockout = *npq4* and overexpression = L17) and VDE (*npq1*) alleles.

Fig. S3: Thylakoid membrane protein analyses of cp26 mutants crossed with contrasting PsbS alleles (knockout = *npq4* and overexpression = L17).

Fig. S4: L17 T-DNA insertion site.

Supplementary data file (statistical analyses)

Supplementary image file (original images)

**Table S1:**
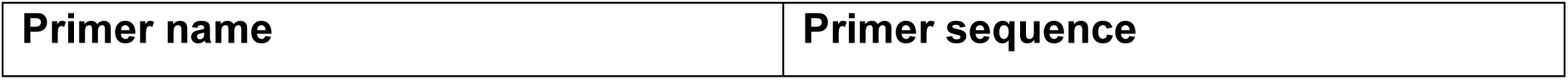
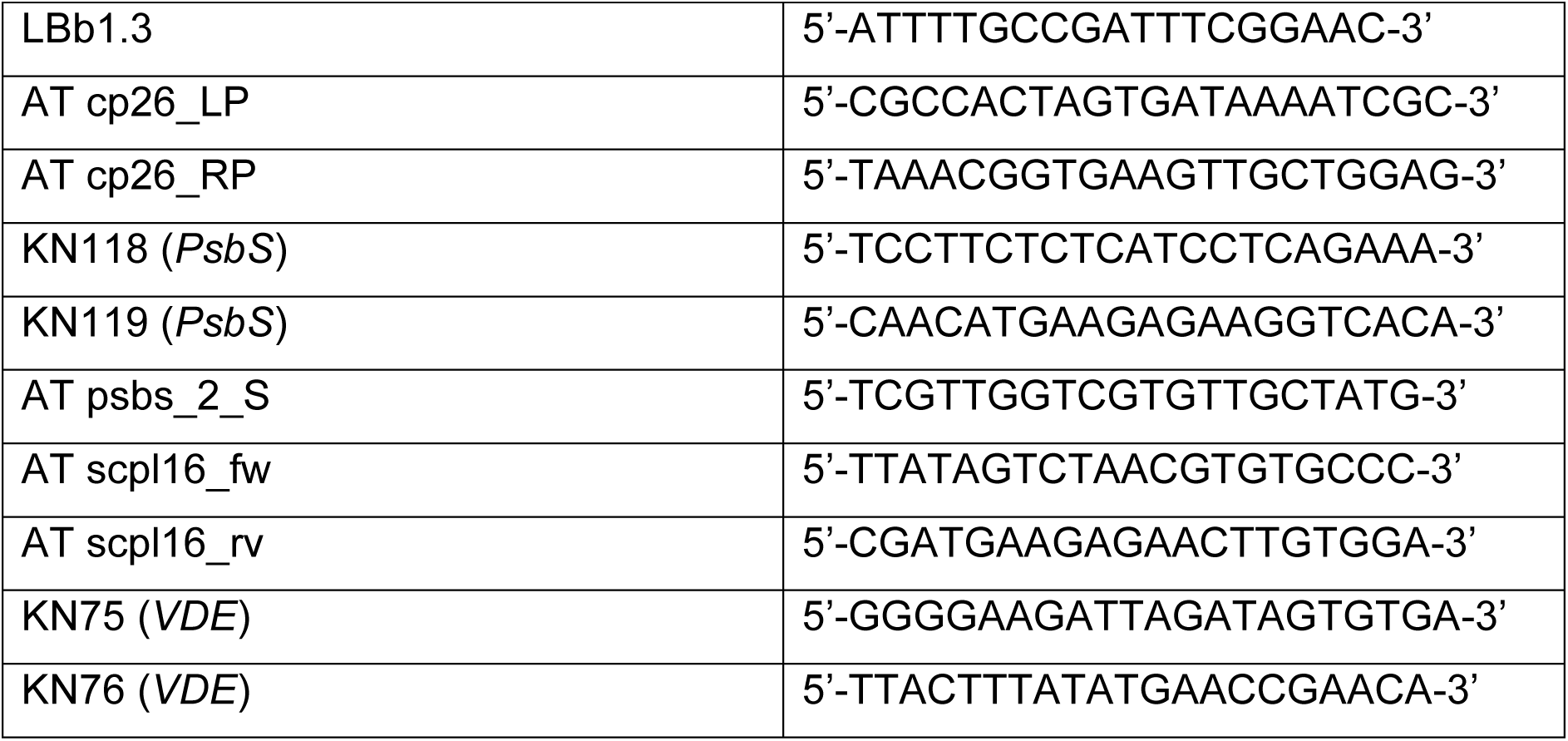
Primers used in this study.

## References

Adams PG, Vasilev C, Hunter CN, Johnson MP. Correlated fluorescence quenching and topographic mapping of Light-Harvesting Complex II within surface-assembled aggregates and lipid bilayers. Biochim Biophys Acta Bioenerg. 2018:1859(10):1075–1085. 10.1016/j.bbabio.2018.06.011

Ahn TK, Avenson TJ, Ballottari M, Cheng YC, Niyogi KK, Bassi R, Fleming GR. Architecture of a charge-transfer state regulating light harvesting in a plant antenna protein. Science. 2008:320(5877):794–797. 10.1126/science.1154800

Andersson J, Walters RG, Horton P, Jansson S. Antisense inhibition of the photosynthetic antenna proteins CP29 and CP26: implications for the mechanism of protective energy dissipation. Plant Cell. 2001:13(5):1193–1204. 10.1105/tpc.13.5.1193

Aro EM, Suorsa M, Rokka A, Allahverdiyeva Y, Paakkarinen V, Saleem A, Battchikova N, Rintamäki E. Dynamics of photosystem II: a proteomic approach to thylakoid protein complexes. J Exp Bot. 2005:56(411):347–356. 10.1093/jxb/eri041

Avenson TJ, Ahn TK, Zigmantas D, Niyogi KK, Li Z, Ballottari M, Bassi R, Fleming GR. Zeaxanthin radical cation formation in minor light-harvesting complexes of higher plant antenna. J Biol Chem. 2008:283(6):3550–3558. 10.1074/jbc.M705645200

Avenson TJ, Ahn TK, Niyogi KK, Ballottari M, Bassi R, Fleming GR. Lutein can act as a switchable charge transfer quencher in the CP26 light-harvesting complex. J Biol Chem. 2009:284(5):2830–2835. 10.1074/jbc.M807192200

Baker NR, Harbinson J, Kramer DM. Determining the limitations and regulation of photosynthetic energy transduction in leaves. Plant Cell Environ. 2007:30(9):1107–1125. 10.1111/j.1365-3040.2007.01680.x

Ballottari M, Mozzo M, Croce R, Morosinotto T, Bassi R. Occupancy and functional architecture of the pigment binding sites of photosystem II antenna complex Lhcb5. J Biol Chem. 2009:284(12):8103–8113. 10.1074/jbc.M808326200

Ballottari M, Mozzo M, Girardon J, Hienerwadel R, Bassi R. Chlorophyll triplet quenching and photoprotection in the higher plant monomeric antenna protein Lhcb5. J Phys Chem B. 2013:117(38):11337–11348. 10.1021/jp402977y

Bassi R, Dall’Osto L. Dissipation of light energy absorbed in excess: the molecular mechanisms. Annu Rev Plant Biol. 2021:72:47–76. 10.1146/annurev-arplant-071720-015522

Bergantino E, Segalla A, Brunetta A, Teardo E, Rigoni F, Giacometti GM, Szabò I. Light-and pH-dependent structural changes in the PsbS subunit of photosystem II. Proc Natl Acad Sci U S A. 2003:100(25):15265–15270. 10.1073/pnas.2533072100

Betterle N, Ballottari M, Zorzan S, de Bianchi S, Cazzaniga S, Dall’Osto L, Morosinotto T, Bassi R. Light-induced dissociation of an antenna hetero-oligomer is needed for non-photochemical quenching induction. J Biol Chem. 2009:284(22):15255–15266. 10.1074/jbc.M808625200

Betterle N, Ballottari M, Baginsky S, Bassi R. High light-dependent phosphorylation of photosystem II inner antenna CP29 in monocots is STN7 independent and enhances nonphotochemical quenching. Plant Physiol. 2015:167(2):457–471. 10.1104/pp.114.252379

Betterle N, Poudyal RS, Rosa A, Wu G, Bassi R, Lee CH. The STN8 kinase-PBCP phosphatase system is responsible for high-light-induced reversible phosphorylation of the PSII inner antenna subunit CP29 in rice. Plant J. 2017:89(4):681–691. 10.1111/tpj.13412

Cazzaniga S, Kim M, Bellamoli F, Jeong J, Lee S, Perozeni F, Pompa A, Jin E, Ballottari M. Photosystem II antenna complexes CP26 and CP29 are essential for nonphotochemical quenching in *Chlamydomonas reinhardtii*. Plant Cell Environ. 2020:43(2):496–509. 10.1111/pce.13680

Cazzaniga S, Kim M, Pivato M, Perozeni F, Sardar S, D’Andrea C, Jin E, Ballottari M. Photosystem II monomeric antenna CP26 plays a key role in nonphotochemical quenching in *Chlamydomonas*. Plant Physiol. 2023:193(2):1365–1380. 10.1093/plphys/kiad391

Chen YE, Ma J, Wu N, Su YQ, Zhang ZW, Yuan M, Zhang HY, Zeng XY, Yuan S. The roles of Arabidopsis proteins of Lhcb4, Lhcb5 and Lhcb6 in oxidative stress under natural light conditions. Plant Physiol Biochem. 2018:130:267–276. 10.1016/j.plaphy.2018.07.014

Cignoni E, Lapillo M, Cupellini L, Acosta-Gutiérrez S, Gervasio FL, Mennucci B. A different perspective for nonphotochemical quenching in plant antenna complexes. Nat Commun. 2021:12(1):7152. 10.1038/s41467-021-27526-8

Correa-Galvis V, Poschmann G, Melzer M, Stühler K, Jahns P. PsbS interactions involved in the activation of energy dissipation in *Arabidopsis*. Nat Plants. 2016:2:15225. 10.1038/nplants.2015.225

Croce R, van Amerongen H. Light-harvesting and structural organization of Photosystem II: from individual complexes to thylakoid membrane. J Photochem Photobiol B. 2011:104(1-2):142–153. 10.1016/j.jphotobiol.2011.02.015

Dall’Osto L, Caffarri S, Bassi R. A mechanism of nonphotochemical energy dissipation, independent from PsbS, revealed by a conformational change in the antenna protein CP26. Plant Cell. 2005:17(4):1217–1232. 10.1105/tpc.104.030601

Dall’Osto L, Cazzaniga S, Bressan M, Paleček D, Židek K, Niyogi KK, Fleming GR, Zigmantas D, Bassi R. Two mechanisms for dissipation of excess light in monomeric and trimeric light-harvesting complexes. Nat Plants. 2017:3:17033. 10.1038/nplants.2017.33

Daskalakis V, Papadatos S, Kleinekathöfer U. Fine tuning of the photosystem II major antenna mobility within the thylakoid membrane of higher plants. Biochim Biophys Acta Biomembr. 2019:1861(12):183059. 10.1016/j.bbamem.2019.183059

de Bianchi S, Dall’Osto L, Tognon G, Morosinotto T, Bassi R. Minor antenna proteins CP24 and CP26 affect the interactions between Photosystem II subunits and the electron transport rate in grana membranes of *Arabidopsis*. Plant Cell. 2008:20(4):1012–1028. 10.1105/tpc.107.055749

de Bianchi S, Betterle N, Kouril R, Cazzaniga S, Boekema E, Bassi R, Dall’Osto L. *Arabidopsis* mutants deleted in the light-harvesting protein Lhcb4 have a disrupted Photosystem II macrostructure and are defective in photoprotection. Plant Cell. 2011:23(7):2659–2679. 10.1105/tpc.111.087320

Dong L, Tu W, Liu K, Sun R, Liu C, Wang K, Yang C. The PsbS protein plays important roles in photosystem II supercomplex remodeling under elevated light conditions. J Plant Physiol. 2015:172:33–41. 10.1016/j.jplph.2014.06.003

Elias E, Liguori N, Croce R. The origin of pigment-binding differences in CP29 and LHCII: the role of protein structure and dynamics. Photochem Photobiol Sci. 2023:22(6):1279–1297. 10.1007/s43630-023-00368-7

Feng X, Pan X, Li M, Pieper J, Chang W, Jankowiak R. Spectroscopic study of the light-harvesting CP29 antenna complex of photosystem II. J Phys Chem B. 2013:117(22):6585–6602. 10.1021/jp4004328

Gacek DA, Holleboom CP, Liao PN, Negretti M, Croce R, Walla PJ. Carotenoid dark state to chlorophyll energy transfer in isolated light-harvesting complexes CP24 and CP29. Photosynth Res. 2020:143(1):19–30. 10.1007/s11120-019-00676-z

Gao S, Zheng Z, Wang J, Wang G. Slow zeaxanthin accumulation and the enhancement of CP26 collectively contribute to an atypical non-photochemical quenching in macroalga *Ulva prolifera* under high light. J Phycol. 2020:56(2):393–403. 10.1111/jpy.12958

Gilmore AM, Hazlett TL, Govindjee. Xanthophyll cycle-dependent quenching of photosystem II chlorophyll a fluorescence: formation of a quenching complex with a short fluorescence lifetime. Proc Natl Acad Sci U S A. 1995:92(6):2273–2277. 10.1073/pnas.92.6.2273

Goral TK, Johnson MP, Duffy CDP, Brain APR, Ruban AV, Mullineaux CW. Light-harvesting antenna composition controls the macrostructure and dynamics of thylakoid membranes in *Arabidopsis*. Plant J. 2012:69(2):289–301. 10.1111/j.1365-313X.2011.04790.x

Guardini Z, Bressan M, Caferri R, Bassi R, Dall’Osto L. Identification of a pigment cluster catalysing fast photoprotective quenching response in CP29. Nat Plants. 2020:6(3):303–313. 10.1038/s41477-020-0612-8

Guardini Z, Gomez RL, Caferri R, Dall’Osto L, Bassi R. Loss of a single chlorophyll in CP29 triggers re-organization of the Photosystem II supramolecular assembly. Biochim Biophys Acta Bioenerg. 2022:1863(5):148555. 10.1016/j.bbabio.2022.148555

Holt NE, Zigmantas D, Valkunas L, Li XP, Niyogi KK, Fleming GR. Carotenoid cation formation and the regulation of photosynthetic light harvesting. Science. 2005:307(5708):433–436. 10.1126/science.1105833

Horton P, Ruban AV, Rees D, Pascal AA, Noctor G, Young AJ. Control of the light-harvesting function of chloroplast membranes by aggregation of the LHCII chlorophyll-protein complex. FEBS Lett. 1991:292(1-2):1–4. 10.1016/0014-5793(91)80819-O

Ilíková I, Ilík P, Opatíková M, Arshad R, Nosek L, Karlický V, Kučerová Z, Roudnický P, Pospíšil P, Lazár D, et al. Towards spruce-type photosystem II: consequences of the loss of light-harvesting proteins LHCB3 and LHCB6 in *Arabidopsis*. Plant Physiol. 2021:187(4):2691–2715. 10.1093/plphys/kiab396

Jansson S. A guide to the Lhc genes and their relatives in *Arabidopsis*. Trends Plant Sci. 1999:4(6):236–240. 10.1016/S1360-1385(99)01419-3

Järvi S, Suorsa M, Paakkarinen V, Aro EM. Optimized native gel systems for separation of thylakoid protein complexes: novel super-and mega-complexes. Biochem J. 2011:439(2):207–214. 10.1042/BJ20102155

Kiss AZ, Ruban AV, Horton P. The PsbS protein controls the organization of the photosystem II antenna in higher plant thylakoid membranes. J Biol Chem. 2008:283(7):3972–3978. 10.1074/jbc.M707410200

Kovács L, Damkjaer J, Kereïche S, Ilioaia C, Ruban AV, Boekema EJ, Jansson S, Horton P. Lack of the light-harvesting complex CP24 affects the structure and function of the grana membranes of higher plant chloroplasts. Plant Cell. 2006:18(11):3106–3120. 10.1105/tpc.106.045641

Krishnan M, Moolenaar GF, Gupta KBSS, Goosen N, Pandit A. Large-scale *in vitro* production, refolding and dimerization of PsbS in different microenvironments. Sci Rep. 2017:7(1):15200. 10.1038/s41598-017-15068-3

Leuenberger M, Morris JM, Chan AM, Leonelli L, Niyogi KK, Fleming GR. Dissecting and modeling zeaxanthin-and lutein-dependent nonphotochemical quenching in *Arabidopsis thaliana*. Proc Natl Acad Sci U S A. 2017:114(33):E7009–E7017. 10.1073/pnas.1704502114

Li XP, Björkman O, Shih C, Grossman AR, Rosenquist M, Jansson S, Niyogi KK. A pigment-binding protein essential for regulation of photosynthetic light harvesting. Nature. 2000:403(6768):391–395. 10.1038/35000131

Li XP, Phippard A, Pasari J, Niyogi KK. Structure-function analysis of photosystem II subunit S (PsbS) *in vivo*. Funct Plant Biol. 2002a:29(10):1131–1139. 10.1071/FP02065

Li XP, Müller-Moulé P, Gilmore AM, Niyogi KK. PsbS-dependent enhancement of feedback de-excitation protects photosystem II from photoinhibition. Proc Natl Acad Sci U S A. 2002b:99(23):15222–15227. 10.1073/pnas.232447699

Li XP, Gilmore AM, Caffarri S, Bassi R, Golan T, Kramer D, Niyogi KK. Regulation of photosynthetic light harvesting involves intrathylakoid lumen pH sensing by the PsbS protein. J Biol Chem. 2004:279(22):22866–22874. 10.1074/jbc.M402461200

Li Z, Ahn TK, Avenson TJ, Ballottari M, Cruz JA, Kramer DM, Bassi R, Fleming GR, Keasling JD, Niyogi KK. Lutein accumulation in the absence of zeaxanthin restores nonphotochemical quenching in the *Arabidopsis thaliana npq1* mutant. Plant Cell. 2009:21(6):1798–1812. 10.1105/tpc.109.066571

Liguori N, Campos SRR, Baptista AM, Croce R. Molecular anatomy of plant photoprotective switches: the sensitivity of PsbS to the environment, residue by residue. J Phys Chem Lett. 2019:10(8):1737–1742. 10.1021/acs.jpclett.9b00437

Marulanda Valencia W, Pandit A. Photosystem II Subunit S (PsbS): a nano regulator of plant photosynthesis. J Mol Biol. 2024:436(5):168407. 10.1016/j.jmb.2023.168407

Mascoli V, Novoderezhkin V, Liguori N, Xu P, Croce R. Design principles of solar light harvesting in plants: functional architecture of the monomeric antenna CP29. Biochim Biophys Acta Bioenerg. 2020:1861(3):148156. 10.1016/j.bbabio.2020.148156

Miloslavina Y, Wehner A, Lambrev PH, Wientjes E, Reus M, Garab G, Croce R, Holzwarth AR. Far-red fluorescence: a direct spectroscopic marker for LHCII oligomer formation in non-photochemical quenching. FEBS Lett. 2008:582(25-26):3625–3631. 10.1016/j.febslet.2008.09.044

Miloslavina Y, de Bianchi S, Dall’Osto L, Bassi R, Holzwarth AR. Quenching in *Arabidopsis thaliana* mutants lacking monomeric antenna proteins of photosystem II. J Biol Chem. 2011:286(42):36830–36840. 10.1074/jbc.M111.273227

Morosinotto T, Baronio R, Bassi R. Dynamics of chromophore binding to Lhc proteins *in vivo* and *in vitro* during operation of the xanthophyll cycle. J Biol Chem. 2002:277(40):36913–36920. 10.1074/jbc.M205339200

Mullineaux CW, Pascal AA, Horton P, Holzwarth AR. Excitation-energy quenching in aggregates of the LHC II chlorophyll-protein complex: a time-resolved fluorescence study. Biochim Biophys Acta Bioenerg. 1993:1141(1):23–28. 10.1016/0005-2728(93)90184-H

Natali A, Gruber JM, Dietzel L, Stuart MCA, van Grondelle R, Croce R. Light-harvesting complexes (LHCs) cluster spontaneously in membrane environment leading to shortening of their excited state lifetimes. J Biol Chem. 2016:291(32):16730–16739. 10.1074/jbc.M116.730101

Nicol L, Nawrocki WJ, Croce R. Disentangling the sites of non-photochemical quenching in vascular plants. Nat Plants. 2019:5(11):1177–1183. 10.1038/s41477-019-0526-5

Nicol L, Croce R. The PsbS protein and low pH are necessary and sufficient to induce quenching in the light-harvesting complex of plants LHCII. Sci Rep. 2021:11(1):7415. 10.1038/s41598-021-86975-9

Nilkens M, Kress E, Lambrev P, Miloslavina Y, Müller M, Holzwarth AR, Jahns P. Identification of a slowly inducible zeaxanthin-dependent component of non-photochemical quenching of chlorophyll fluorescence generated under steady-state conditions in *Arabidopsis*. Biochim Biophys Acta Bioenerg. 2010:1797(4):466–475. 10.1016/j.bbabio.2010.01.001

Niyogi KK, Grossman AR, Björkman O. *Arabidopsis* mutants define a central role for the xanthophyll cycle in the regulation of photosynthetic energy conversion. Plant Cell. 1998:10(7):1121–1134. 10.1105/tpc.10.7.1121

Pashayeva A, Wu G, Huseynova I, Lee CH, Zulfugarov IS. Role of thylakoid protein phosphorylation in energy-dependent quenching of chlorophyll fluorescence in rice plants. Int J Mol Sci. 2021:22(15):7978. 10.3390/ijms22157978

Pawlak K, Paul S, Liu C, Reus M, Yang C, Holzwarth AR. On the PsbS-induced quenching in the plant major light-harvesting complex LHCII studied in proteoliposomes. Photosynth Res. 2020:144(2):195–208. 10.1007/s11120-020-00740-z

Peers G, Truong TB, Ostendorf E, Busch A, Elrad D, Grossman AR, Hippler M, Niyogi KK. An ancient light-harvesting protein is critical for the regulation of algal photosynthesis. Nature. 2009:462(7272):518–521. 10.1038/nature08587

Peng X, Deng X, Tang X, Tan T, Zhang D, Liu B, Lin H. Involvement of Lhcb6 and Lhcb5 in photosynthesis regulation in *Physcomitrella patens* response to abiotic stress. Int J Mol Sci. 2019:20(15):3665. 10.3390/ijms20153665

Pinnola A, Staleva-Musto H, Capaldi S, Ballottari M, Bassi R, Polívka T. Electron transfer between carotenoid and chlorophyll contributes to quenching in the LHCSR1 protein from *Physcomitrella patens*. Biochim Biophys Acta Bioenerg. 2016:1857(12):1870–1878. 10.1016/j.bbabio.2016.09.001

Porra RJ, Thompson WA, Kriedemann PE. Determination of accurate extinction coefficients and simultaneous equations for assaying chlorophylls *a* and *b* extracted with four different solvents: verification of the concentration of chlorophyll standards by atomic absorption spectroscopy. Biochim Biophys Acta Bioenerg. 1989:975(3):384–394. 10.1016/S0005-2728(89)80347-0

Ruban AV, Walters RG, Horton P. The molecular mechanism of the control of excitation energy dissipation in chloroplast membranes: inhibition of delta pH-dependent quenching of chlorophyll fluorescence by dicyclohexylcarbodiimide. FEBS Lett. 1992:309(2):175–179. 10.1016/0014-5793(92)81089-5

Ruban AV, Wilson S. The mechanism of non-photochemical quenching in plants: localization and driving forces. Plant Cell Physiol. 2021:62(7):1063–1072. 10.1093/pcp/pcaa155

Saccon F, Giovagnetti V, Shukla MK, Ruban AV. Rapid regulation of photosynthetic light harvesting in the absence of minor antenna and reaction centre complexes. J Exp Bot. 2020:71(12):3626–3637. 10.1093/jxb/eraa126

Sacharz J, Giovagnetti V, Ungerer P, Mastroianni G, Ruban AV. The xanthophyll cycle affects reversible interactions between PsbS and light-harvesting complex II to control non-photochemical quenching. Nat Plants. 2017:3:16225. 10.1038/nplants.2016.225

Sardar S, Caferri R, Camargo FVA, Pamos Serrano J, Ghezzi A, Capaldi S, Dall’Osto L, Bassi R, D’Andrea C, Cerullo G. Molecular mechanisms of light harvesting in the minor antenna CP29 in near-native membrane lipidic environment. J Chem Phys. 2022:156(20):205101. 10.1063/5.0087898

Sardar S, Caferri R, Camargo FVA, Capaldi S, Ghezzi A, Dall’Osto L, D’Andrea C, Cerullo G, Bassi R. Site-directed mutagenesis of the chlorophyll-binding sites modulates excited-state lifetime and chlorophyll-xanthophyll energy transfer in the monomeric light-harvesting complex CP29. J Phys Chem Lett. 2024:15(11):3149–3158. 10.1021/acs.jpclett.3c02900

Sárvári É, Gellén G, Sági-Kazár M, Schlosser G, Solymosi K, Solti Á. Qualitative and quantitative evaluation of thylakoid complexes separated by Blue Native PAGE. Plant Methods. 2022:18(1):23. 10.1186/s13007-022-00858-2

Shukla MK, Watanabe A, Wilson S, Giovagnetti V, Moustafa EI, Minagawa J, Ruban AV. A novel method produces native light-harvesting complex II aggregates from the photosynthetic membrane revealing their role in nonphotochemical quenching. J Biol Chem. 2020:295(51):17816–17826. 10.1074/jbc.RA120.016181

Singer T, Burke E. High-throughput TAIL-PCR as a tool to identify DNA flanking insertions. Methods Mol Biol. 2003:236:241–272. 10.1385/1-59259-413-1:241

Steen CJ, Morris JM, Short AH, Niyogi KK, Fleming GR. Complex roles of PsbS and xanthophylls in the regulation of nonphotochemical quenching in *Arabidopsis thaliana* under fluctuating light. J Phys Chem B. 2020:124(46):10311–10325. 10.1021/acs.jpcb.0c06265

Suorsa M, Rantala M, Mamedov F, Lespinasse M, Trotta A, Grieco M, Vuorio E, Tikkanen M, Järvi S, Aro EM. Light acclimation involves dynamic re-organization of the pigment-protein megacomplexes in non-appressed thylakoid domains. Plant J. 2015:84(2):360–373. 10.1111/tpj.13004

Sylak-Glassman EJ, Zaks J, Amarnath K, Leuenberger M, Fleming GR. Characterizing non-photochemical quenching in leaves through fluorescence lifetime snapshots. Photosynth Res. 2016:127(1):69–76. 10.1007/s11120-015-0104-2

Tian L, Dinc E, Croce R. LHCII populations in different quenching states are present in the thylakoid membranes in a ratio that depends on the light conditions. J Phys Chem Lett. 2015:6(12):2339–2344. 10.1021/acs.jpclett.5b00944

Tietz S, Hall CC, Cruz JA, Kramer DM. NPQ(T): a chlorophyll fluorescence parameter for rapid estimation and imaging of non-photochemical quenching of excitons in photosystem-II-associated antenna complexes. Plant Cell Environ. 2017:40(8):1243–1255. 10.1111/pce.12924

Townsend AJ, Saccon F, Giovagnetti V, Wilson S, Ungerer P, Ruban AV. The causes of altered chlorophyll fluorescence quenching induction in the *Arabidopsis* mutant lacking all minor antenna complexes. Biochim Biophys Acta Bioenerg. 2018:1859(9):666–675. 10.1016/j.bbabio.2018.03.005

Tutkus M, Chmeliov J, Trinkunas G, Akhtar P, Lambrev PH, Valkunas L. Aggregation-related quenching of LHCII fluorescence in liposomes revealed by single-molecule spectroscopy. J Photochem Photobiol B. 2021:218:112174. 10.1016/j.jphotobiol.2021.112174

Walters RG, Ruban AV, Horton P. Identification of proton-active residues in a higher plant light-harvesting complex. Proc Natl Acad Sci U S A. 1996:93(24):14204–14209. 10.1073/pnas.93.24.14204

Ware MA, Giovagnetti V, Belgio E, Ruban AV. PsbS protein modulates non-photochemical chlorophyll fluorescence quenching in membranes depleted of photosystems. J Photochem Photobiol B. 2015:152(Part B):301–307. 10.1016/j.jphotobiol.2015.07.016

Wilk L, Grunwald M, Liao P-N, Walla PJ, Kühlbrandt W. Direct interaction of the major light-harvesting complex II and PsbS in nonphotochemical quenching. Proc Natl Acad Sci U S A. 2013:110(14):5452–5456. 10.1073/pnas.1205561110

Wilson S, Li DH, Ruban AV. The structural and spectral features of light-harvesting complex II proteoliposomes mimic those of native thylakoid membranes. J Phys Chem Lett. 2022:13(24):5683–5691. 10.1021/acs.jpclett.2c01019

Xu P, Tian L, Kloz M, Croce R. Molecular insights into zeaxanthin-dependent quenching in higher plants. Sci Rep. 2015:5:13679. 10.1038/srep13679

Yakushevska AE, Keegstra W, Boekema EJ, Dekker JP, Andersson J, Jansson S, Ruban AV, Horton P. The structure of photosystem II in *Arabidopsis*: localization of the CP26 and CP29 antenna complexes. Biochemistry. 2003:42(3):608–613. 10.1021/bi027109z

